# An *in silico* comparison of reduced-representation and sequence-capture protocols for phylogenomics

**DOI:** 10.1101/032565

**Authors:** Rupert A. Collins, Tomas Hrbek

## Abstract

In the age of genome-scale DNA sequencing, choice of molecular marker arguably remains an important decision in planning a phylogenetic study. Using published genomes from 23 primate species, we make a standardized comparison of four of the most frequently used protocols in phylogenomics, *viz*., targeted sequence-enrichment using ultraconserved element and exon-capture probes, and reduced genomic representation using restriction-site-associated DNA sequencing (RADseq and ddRAD-seq). Here we present a procedure to perform *in silico* extractions from genomes and create directly comparable datasets for each class of marker. We then compare these datasets in terms of both phylogenetic resolution and ability to consistently and precisely estimate clade ages using fossil-calibrated molecular-clock models. Furthermore, we were also able to directly compare these results to previously published datasets from Sanger-sequenced nuclear exons and mitochondrial genomes under the same analytical conditions. Our results show—although with the exception of the mitochondrial genome and ddRADseq datasets—that for uncontroversial nodes all data classes performed equally well, i.e. they recovered the same well supported topology. However, for one difficult-to-resolve node comprising a rapid diversification (subfamilial relationships among the Cebidae), we report well supported but conflicting topologies among the marker classes, likely the result of mismodelling of gene tree heterogeneity. Likewise, clade age estimates showed consistent discrepancies between datasets; for recent nodes, clade ages estimated by nuclear exon datasets were younger than those of the UCE, RAD and mitochondrial data, but *vice versa* for the deepest nodes in the primate phylogeny. This effect can be explained by temporal differences in phylogenetic informativeness and choice of clock model used. Finally, we conclude by emphasizing that while huge numbers of loci are probably not required for uncontroversial phylogenetic questions—for which practical considerations such as cost and ease of data generation/sharing/aggregating therefore become increasingly important—accurately modelling heterogeneous data remains as relevant as ever for the more recalcitrant problems.

## Introduction

The development of high-throughput DNA sequencing technologies initiated a gradual move away from the traditional amplicon-based Sanger-sequencing method for generating data for molecular phylogenetics (McCormack *et al*., 2013; Lemmon & Lemmon, 2013). Researchers are now no longer restricted to using a small number of well known and often group-specific loci, and at the same time can generate great volumes of genome-wide data at a substantially lower per-base cost. There is also now longer a requirement for significant *a priori* knowledge of the genome, and so data can readily be collected from non-model organisms (Davey & Blaxter, 2010; Lemmon *et al*., 2012). However, until whole genome sequencing is financially viable for phylogenetic projects with dense/broad taxon coverage, there still remains the need for an *a priori* choice of which genomic regions are to be sequenced and used in phylogenetic analyses. Two general methodologies have been the most widely used in empirical phylogenomic studies: “reduced representation” and “sequence capture” (also referred to as “targeted enrichment”). The aim of this study is to make a standardized comparison of the two methods in terms of their phylogenetic performance.

### Reduced genomic representation

Reduced representations of a genome can be created by enzymatic digestion of template DNAs at restriction sites, followed by adapter ligation, size selection and the sequencing of the restriction-enzyme-associated flanking sites on a high-throughput platform. This technique is most commonly known as restriction-site-associated DNA sequencing, or “RADseq” (Baird *et al*., 2008; Davey *et al*., 2011). Several different RADseq protocols exist, for example double-digest RADseq (ddRADseq), and each has its own pros and cons (Puritz *et al*., 2014). Although initially developed for population genetic studies, RADseq data have been applied to interspecific phylogenetic problems, particularly in non-vertebrate groups (e.g. Cruaud *et al*., 2014; Eaton & Ree, 2013; Rubin *et al*., 2012). One of the key issues arising from these studies is whether a sufficient number of restriction sites are conserved across increasingly divergent species to permit resolution of the deeper nodes of a tree (Cariou *et al*., 2013; Rubin *et al*., 2012).

### Targeted enrichment and sequence capture

An alternative approach to accessing phylogenetically representative genomic regions is via the use of targeted enrichment, whereby templates are hybridized to oligonucleotide probes and sequenced along with their flanking sites (Faircloth *et al*., 2012b; Lemmon *et al*., 2012). Although *a priori* knowledge of the genomes are required to design the probes, sufficient genomic resources are now available for many groups to make this feasible (Bragg *et al*., 2015).

One such target of targeted enrichment are the ultraconserved elements (UCEs), highly conserved but probably functional sections of non-coding DNA scattered across the genome (Katzman *et al*., 2007). While the UCE cores themselves can be informative, it is their flanking regions that contain the majority of the phylogenetic information, and this information increases with distance from the UCE core (Gilbert *et al*., 2015; Smith *et al*., 2014). Faircloth *et al*. (2012b) provided the first protocol and probeset for amniotes comprising 2,386 UCEs, and this was followed by a probeset for fishes (Faircloth *et al*., 2013). While UCEs have been used mainly for deep phylogenetic studies (e.g. Crawford *et al*., 2012; Faircloth *et al*., 2013; McCormack *et al*., 2012a,b), they have also proved useful in studies of recently diverged species such as cis- and trans-Andean bird species pairs (Harvey *et al*., 2013; Smith *et al*., 2014).

“Anchored hybrid enrichment” is another targeted enrichment method. It targets exons and their flanking regions, but in contrast to the UCEs, most of the phylogenetic information comes from the targeted element, the exons. A 512-locus probeset has been developed by Lemmon *et al*. (2012), and the protocol has been used to generate data for phylogenetic studies of snakes (Pyron *et al*., 2014), lizards (Leaché *et al*., 2014) and fishes (Eytan *et al*., 2015). Although this technique captures more than exon data—the proportion of intronic data increases as distance from the probes increases (see Fig. 2 of Lemmon *et al*., 2012), however, the final loci are predominantly coding regions (around 80%)—for brevity we refer to the method as “exon capture (exon-cap)” from herein.

### Study aims

Obtaining a set of loci representative of the phylogenetic history of the organisms under study is the primary motivation when choosing a protocol, but practical and technical issues are also important in ensuring experimental data are collected in a cost effective manner. Therefore when using high-throughput sequencing platforms it is therefore important to ask *a priori* how much data is sufficient to answer the question with confidence, and therefore avoid unnecessarily sequencing an excessive number of loci/sites at the expense of the number of individuals or the sequencing depth.

Here, we aim to provide an assessment of four published protocols: RADseq using the *SdaI* enzyme (Baird *et al*., 2008); ddRADseq using the *SdaI* and *Csp6I* enzymes (Peterson *et al*., 2012); UCEs using the 2,386-locus probeset of Faircloth *et al*. (2012b); and exon-cap using the 512-locus probeset of Lemmon *et al*. (2012). We aim to examine the ability of the data generated using these protocols to resolve a “known” phylogeny and provide precise clade-age estimates. Specifically, for each dataset we will measure: (1) the number of recovered loci and proportions of missing data; (2) topological uncertainty and statistical support for resolving dichotomies; (3) consistency in branch length estimates after application of fossil-calibrated molecular-clock models; and (4) phylogenetic informativeness over a standardized time scale.

To facilitate this *in silico* study we will use 23 primate species that have whole-genome data available. Primates were chosen since complete genomes became available in GenBank at the end of March 2015 and thus could be mined for the different data types we wanted to compare. Secondly, given that comprehensive phylogenies of primates have already been published (Finstermeier *et al*., 2013; Jameson *et al*., 2011; Perelman *et al*., 2011)—with the backbone primate phylogeny largely resolved and only a small number of controversial relationships remaining (e.g. the cebid subfamilies; Wildman *et al*., 2009)—we can therefore compare our results directly to those of Perelman *et al*. (2011, Sanger-sequenced nuclear exons) and Finstermeier *et al*. (2013, mitochondrial genomes) under identical analytical conditions.

## Materials and methods

All data, scripts and program settings to repeat this work are available^1^ from a public git repository hosted at https://github.com/boopsboops/primates-genome-phylo.

### Genomic data extraction

Complete genomes of 23 primate species were downloaded from NCBI GenBank (http://www.ncbi.nlm.nih.gov/genome) in April 2015. A complete list is provided in Table 1. *Daubentonia madagascariensis* was not included since the genome was not assembled into scaffolds at the time of writing. *Eulemur flavifrons* and *Eulemur macaco* genomes were not released until August 2015, and thus were not included in this study either.

**Table 1.**
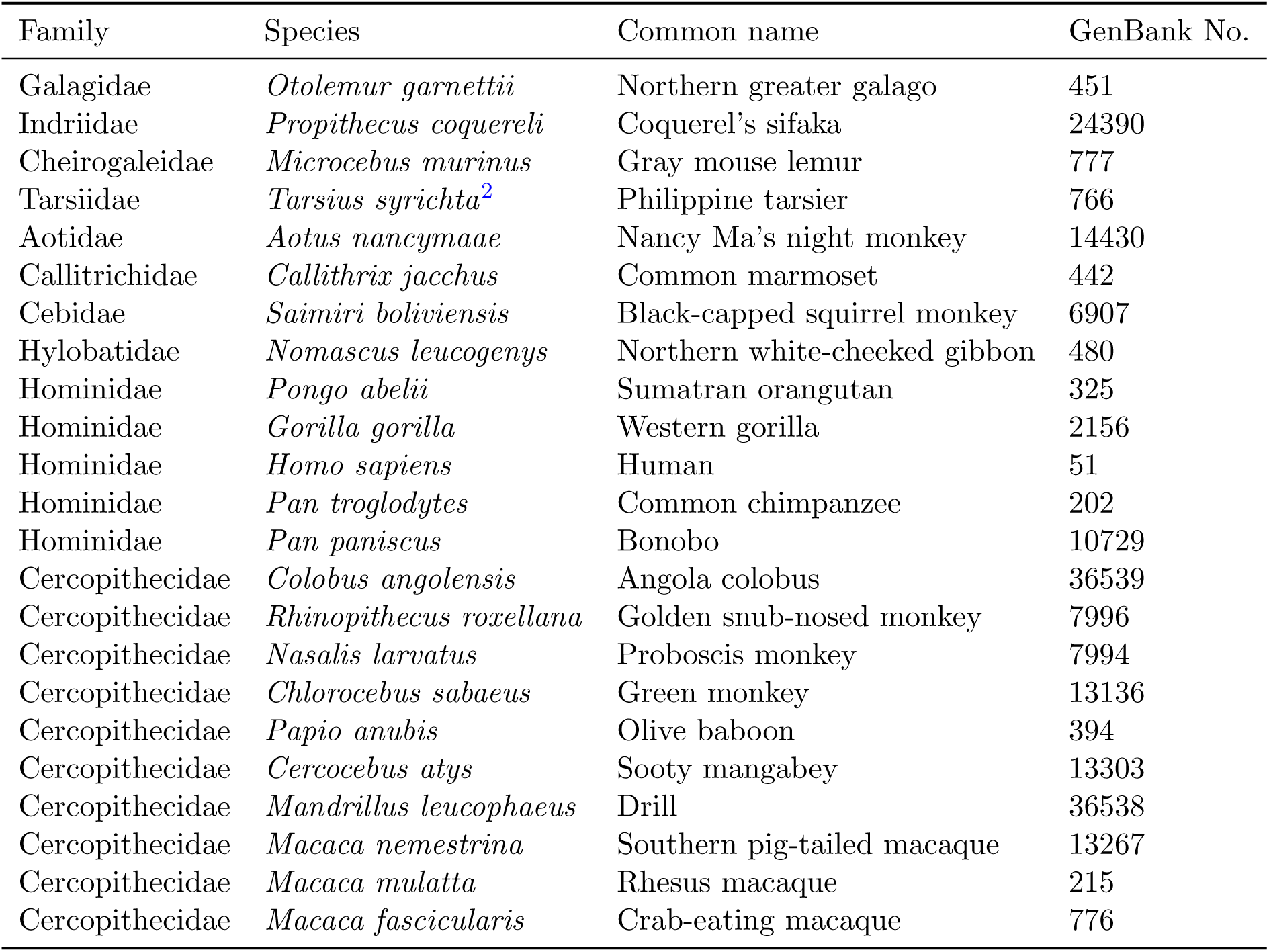
List of species and GenBank genome accession numbers. Resources can be accessed at http://www.ncbi.nlm.nih.gov/genome/[insertGenBankNo.].^2^

| Family | Species | Common name | GenBank No. |
| --- | --- | --- | --- |
| Galagidae | *Otolemur garnettii* | Northern greater galago | 451 |
| Indridae | *Propithecus coquereli* | Coquerel’s sifaka | 24390 |
| Cheirogaleidae | *Microcebus murinus* | Gray mouse lemur | 777 |
| Tarsiidae | *Tarsius syrichta*<sup>2</sup> | Philippine tarsier | 766 |
| Aotidae | *Aotus nancymae* | Nancy Ma’s night monkey | 14430 |
| Callitrichidae | *Callithrix jacchus* | Common marmoset | 442 |
| Cebidae | *Saimiri boliviensis* | Black-capped squirrel monkey | 6907 |
| Hylobatidae | *Nomascus leucogenys* | Northern white-cheeked gibbon | 480 |
| Hominidae | *Pongo abelii* | Sumatran orangutan | 325 |
| Hominidae | *Gorilla gorilla* | Western gorilla | 2156 |
| Hominidae | *Homo sapiens* | Human | 51 |
| Hominidae | *Pan troglodytes* | Common chimpanzee | 202 |
| Hominidae | *Pan paniscus* | Bonobo | 10729 |
| Cercopithecidae | *Colobus angolensis* | Angola colobus | 36539 |
| Cercopithecidae | *Rhinopithecus roxellana* | Golden snub-nosed monkey | 7996 |
| Cercopithecidae | *Nasalis larvatus* | Proboscis monkey | 7994 |
| Cercopithecidae | *Chlorocebus sabaeus* | Green monkey | 13136 |
| Cercopithecidae | *Papio anubis* | Olive baboon | 394 |
| Cercopithecidae | *Cercocebus atys* | Sooty mangabey | 13303 |
| Cercopithecidae | *Mandrillus leucophaeus* | Drill | 36538 |
| Cercopithecidae | *Macaca nemestrina* | Southern pig-tailed macaque | 13267 |
| Cercopithecidae | *Macaca mulatta* | Rhesus macaque | 215 |
| Cercopithecidae | *Macaca fascicularis* | Crab-eating macaque | 776 |

RADseq data were extracted by performing an *in silico* digest with the *SdaI* restriction enzyme, an 8-cutter enzyme, and saving 300 bp on either side of the restriction cut site. Generated fragments on either side of the restriction enzyme were treated as being unlinked (*sensu* the premises of the RADseq protocol; Baird *et al*., 2008). We implicitly assumed that these data would have been generated on an Illumina HiSeq 2500, and thus we assumed 300 bp paired-end read lengths, 150 bp in each direction.

The ddRADseq data were extracted by performing an *in silico* digest with the *SdaI* restriction enzyme, followed by a digest with the *Csp6I* restriction enzyme, a 6-cutter enzyme. Subsequently all fragments with an *SdaI* cut on one end, an *Csp6I* cut on the other end, and within a 320–400 bp range, were saved. The 320–400 bp range corresponds to the “tight” cut on the PippinPrep centered on 360 bp. The decision to this size range was based on several considerations: (1) the ability to select size ranges no smaller than 80 bp at 300 bp or larger on the PippinPrep; (2) most Illumina paired-end RADseq protocols work with library inserts of 300–500 bp; and (3) the ability of the IonTorrent PGM to sequence fragments of up to 400 bp.

To extract the UCE and exon-cap data, we converted the downloaded genomes into local Blast databases. Given that the 2,560 tiled UCE baits target 2,386 UCEs using 120-mer tiled DNA probes (Faircloth *et al*., 2012b), and the 56,664 tiled exon-cap probes target 512 loci (Lemmon *et al*., 2012), we merged all tiled probes to constituent loci. We subsequently queried these merged probes using MegaBlast against the Blast database of each species, and extracted the probes plus associated flanking regions on either side of the probes. For the UCEs we selected 240 bp flanking either side of probe, and exon-cap we selected 180 bp, in addition to the core probe region. Due to the difference in probe lengths, this resulted in a standardized locus length of 600 bp for both methods.

### Post-processing

Post extraction from the genomes, all output was formatted so that it could be processed in the PyRAD v3.0.5 pipeline (Eaton, 2014). We used PyRAD since it was designed to assemble data for phylogenetic studies where variation between divergent taxa may include indels. PyRAD was used only to cluster loci across species, align the loci, filter for paralogs, assemble the loci into a concatenated matrix, and generate summary statistics (i.e. only the pipeline steps six to seven of PyRAD were used). We formed data matrices for analyses such that at least four individuals shared any given locus, which is the minimum number of individuals needed to form a quartet. Clustering threshold was set to 88% similarity, the default in PyRAD.

Next, each dataset was filtered to keep only the loci that included sequences for at least 12 of the 23 taxa (≈ 50%) using Gblocks (Castresana, 2000); this step also removed any ambiguously aligned sites from the matrices.

### Additional datasets

The Perelman *et al*. (2011) (= “Perelman-exon”) and Finstermeier *et al*. (2013) (= “Finstermeier-mtDNA”) alignments were accessed from online appendices of each study. Because some of the taxa in these studies did not match those of our genomes, some individuals were substituted: for Perelman-exon, *Cercocebus torquatus* replaced *Cercocebus atys* and *Rhinopithecus brelichi* replaced *Rhinopithecus roxellana*; for Finstermeier-mtDNA, *Otolemur crassicaudatus, Cheirogaleus medius, Aotus azarae, Callichtrix geoffroyi, Colobus guereza, Chlorocebus pygerythrus, Mandrillus sphinx, Cercocebus chrysogaster* and *Macaca sylvanus* replaced *Otolemur garnettii, Microcebus murinus, Aotus nancymaae, Callichtrix jacchus, Colobus angolensis, Chlorocebus sabaeus, Mandrillus leucophaeus, Cercocebus atys* and *Macaca nemestrina*, respectively. Both of these datasets were also subjected to the same alignment cleaning step using Gblocks.

### Phylogenetic analyses

Topological uncertainty was assessed using Bayesian posterior probabilities and bipartition data generated by the program ExaBayes (Aberer *et al*., 2014). Two independent runs were carried out from random starting topologies for three million generations for each dataset. Convergence and stationarity of parameter estimates was verified using Tracer (Rambaut *et al*., 2013). Runs were combined and then summarized into maximum clade credibility (MCC) trees using TreeAnnotator (Rambaut & Drummond, 2015) with a 10% burn-in.

### Divergence time analyses

We estimated divergence times using both the strict and uncorrelated (relaxed) molecular clock methods employed in the *chronos* function of the R package Ape (Paradis *et al*., 2004; Paradis, 2013; Sanderson, 2002), and we compared these models using Φ*IC* criterion (Paradis, 2013). To get confidence intervals for each dataset, we dated all 9,000 post-burnin ExaBayes trees. Temporal information was incorporated in the form of six fossil calibration points taken from Perelman *et al*. (2011): root (all primates), B (Simiiformes), C (Catarrhini), E (Papionini), G (Hominidae), and H (*Homo-Pan*); these calibration points are illustrated in Figure 1, and are described in more detail by (Perelman *et al*., 2011). Because *chronos* does not support normally distributed calibration densities, these were converted to uniform distributions by taking the 95th percentile of the normal distribution.

**Figure 1.**
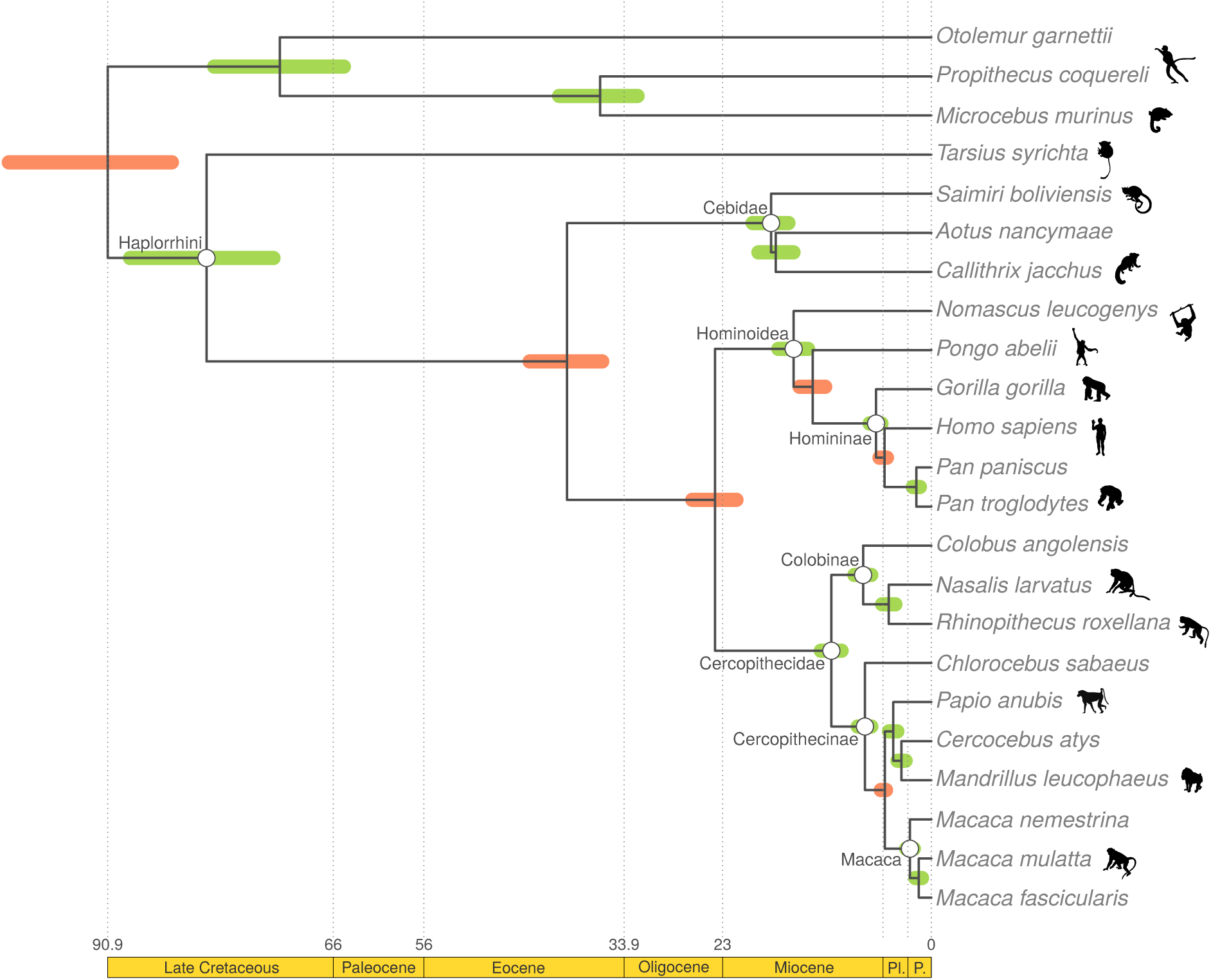
Exemplar phylogenetic tree to show taxon relationships, calibrated nodes, and nodes of interest. Dataset comprises the Gblooks-pruned Perelman-exon dataset; the tree was constructed from 9,000 post-burnin ExaBayes trees dated using the strict clock model in Ape. Calibrated nodes are highlighted in red and follow six calibration points—root (all primates), B (Simiiformes), C (Catarrhini), E (Papionini), G (Hominidae), and H (*Homo-Pan*)—from Perelman *et al*. (2011), while nodes of interest presented in Figure 5 and Figure 6 are indicated with an open circle and clade names. Examples of taxa are also shown (images from http://phylopic.org/).

### Visualization and phylogenetic informativeness

Topological uncertainty in the posterior samples was visualized using RWTY (https://github.com/danlwarren/RWTY) software implemented in R, but based on the “Are We There Yet?” (AWTY) program of Nylander *et al*. (2008). An MCC tree from the strictclock-dated Perelman-exon dataset was used to assess the temporal phylogenetic informativeness (PI) of each dataset using the software Tapir (Faircloth *et al*., 2012a; Townsend, 2007).

## Results

### Summary statistics

The number of recovered loci from the *in silico* bioinformatic extraction steps differed by several orders of magnitude from between 409 (exon-cap) to 235,074 (RADseq), as did the overall number of sites in each dataset (Table 2). After cleaning in Gblocks to remove ambiguously aligned sites and all loci with less than 12 taxa, the size of the datasets (total number of sites) were reduced, often substantially, from their original sizes (Table 3): 93.7% (RADseq), 98.0% (ddRADseq), 61.8% (UCE), and 29.4% (exon-cap).

**Table 2.** Characteristics of the datasets after running through the PyRAD pipeline, and requiring that at least four individuals share a given locus (minimum number of individuals needed to form a quartet).

| Dataset | Number loci | Number sites | No. var. sites | No. parsim. inform. sites |
| --- | --- | --- | --- | --- |
| RAD | 235,074 | 20,774,455 | 2,049,118 | 707,221 |
| ddRAD | 15,913 | 6,202,215 | 456,302 | 145,919 |
| UCE | 2,771 | 1,647,056 | 107,185 | 33,989 |
| Exon-cap | 409 | 238,806 | 26,304 | 11,673 |

**Table 3.** Characteristics of the datasets post cleaning by Gblocks to remove ambiguously aligned sites and limit missing data at the per-locus level to ≈ 50% (loci included if available for at least 12 of 23 taxa).

| Dataset | Number loci | Number sites | No. parsim. inform. sites | Prop. parsim. inform. sites | Prop. miss. data |
| --- | --- | --- | --- | --- | --- |
| RAD | 14,996 | 1,299,570 | 83,530 | 0.064 | 0.44 |
| ddRAD | 329 | 123,768 | 6,977 | 0.056 | 0.45 |
| UCE | 1080 | 629,793 | 19,460 | 0.031 | 0.27 |
| Exon-cap | 290 | 168,682 | 9,861 | 0.058 | 0.18 |
| Perelman-exon | 54 | 31,317 | 3,189 | 0.102 | 0.10 |
| Finstermeier-mtDNA | 37 | 15,059 | 7,014 | 0.466 | 0 |

For all datasets the number of recovered loci declined as the number of taxa increased, although this dropout was substantially more pronounced in the reduced representation datasets (RADseq, ddRADseq) than in the sequence capture (exon-cap, UCE) datasets (Figure 2). The proportion of pairwise shared loci decreased with phylogenetic divergence (Figure 3, Table 4), and followed an exponential function in the RADseq (Δ*AIC_i_* = 110.6; *R*^2^ = 0.81); ddRADseq (Δ*AIC_i_* = 203.3; *R*^2^ = 0.82) and UCE (Δ*AIC_i_* = 299.8; *R*^2^ = 0.95) datasets, but a linear function in the exon-cap dataset (Δ*AIC_i_* = 131.4; *R*^2^ = 0.89).

**Figure 2.**
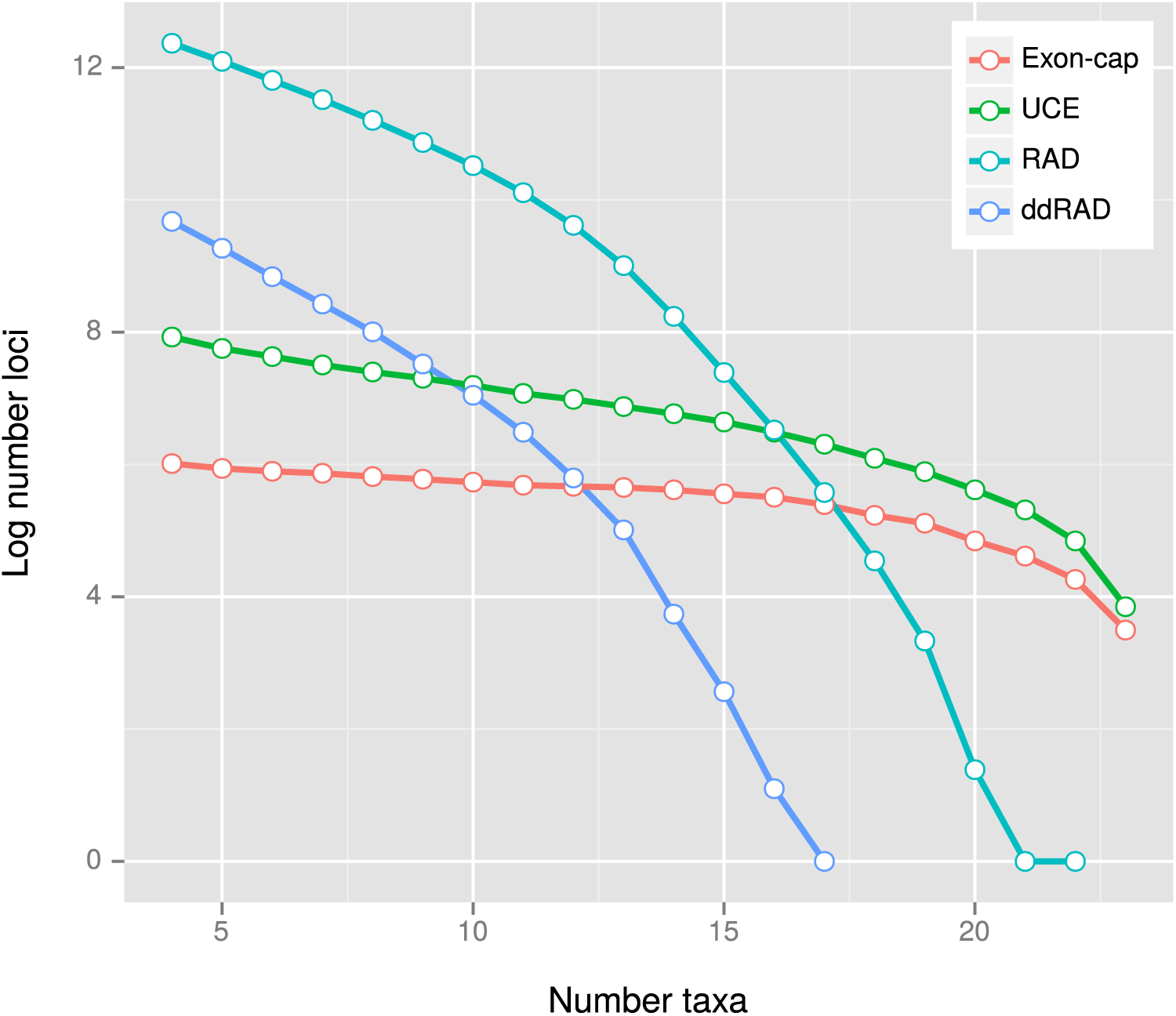
PyRAD pipeline results for the four genome-scale datasets, showing relative dropout of loci as taxa are added, i.e. number of loci (log transformed) for which at least *n* species have data.

**Figure 3.**
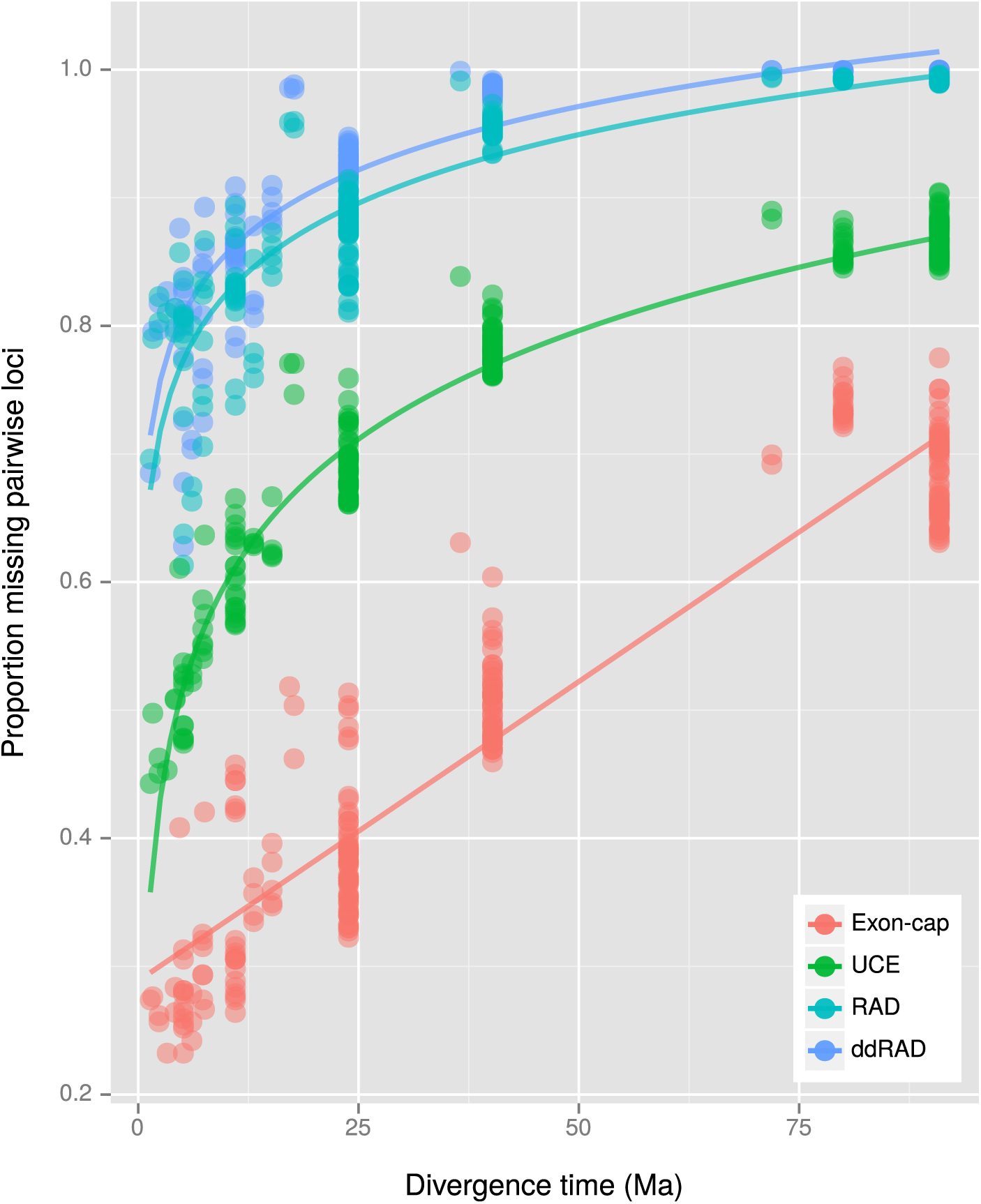
Proportion of missing loci as a function of divergence time for each dataset. The strict clock tree resulting from the analysis of the Perelman-exon dataset was used for the divergence time calculations. Note that the proportion of missing data increases exponentially with divergence time in the RADseq, ddRADseq and UCE datasets, while it increases linearly in the exon-cap dataset.

**Table 4.** Number of loci contributed to the final data matrix by each species, assuming that at least 12 individuals share a given locus (i.e. equivalent to a maximum of ≈ 50% missing data).

| Species | RAD | ddRAD | UCE | Exon-cap |
| --- | --- | --- | --- | --- |
| *Otolemur garnettii* | 1,148 | 6 | 325 | 113 |
| *Propithecus coquereli* | 1,699 | 15 | 394 | 137 |
| *Microcebus murinus* | 1,582 | 11 | 391 | 134 |
| *Tarsius syrichta* | 1,365 | 9 | 401 | 108 |
| *Aotus nancymaae* | 4,212 | 60 | 619 | 180 |
| *Callithrix jacchus* | 5,426 | 63 | 581 | 164 |
| *Saimiri boliviensis* | 3,543 | 44 | 624 | 176 |
| *Nomascus leucogenys* | 7,439 | 166 | 855 | 208 |
| *Pongo abelii* | 12,324 | 266 | 802 | 208 |
| *Gorilla gorilla* | 13,122 | 289 | 916 | 220 |
| *Homo sapiens* | 13,960 | 308 | 918 | 220 |
| *Pan troglodytes* | 13,322 | 291 | 920 | 208 |
| *Pan paniscus* | 8,190 | 199 | 925 | 219 |
| *Colobus angolensis* | 8,561 | 204 | 935 | 225 |
| *Rhinopithecus roxellana* | 8,898 | 195 | 394 | 137 |
| *Nasalis larvatus* | 11,586 | 263 | 802 | 191 |
| *Chlorocebus sabaeus* | 11,769 | 272 | 930 | 216 |
| *Papio anubis* | 13,723 | 300 | 915 | 219 |
| *Cercocebus atys* | 8,962 | 204 | 1,009 | 232 |
| *Mandrillus leucophaeus* | 9,008 | 189 | 1,022 | 230 |
| *Macaca nemestrina* | 7,788 | 196 | 1,006 | 231 |
| *Macaca mulatta* | 14,053 | 311 | 985 | 226 |
| *Macaca fascicularis* | 12,902 | 296 | 1,015 | 223 |

### Phylogenetic performance

Individual phylogenies from the ExaBayes analyses are presented in Supplementary Figure 1 to Supplementary Figure 6. Topologies inferred from all the datasets were sup-ported with high Bayesian posterior probability (BPP (≥ 0.99), except with the following exceptions: (1) the Perelman-exon dataset and the ddRADseq dataset failed to resolve the relationships within the Cebidae; and (2) the ddRADseq dataset failed to resolve the basal relationships at the root of the primates.

The relationships inferred from all the datasets were consistent, but with the following exceptions: (1) the RADseq, ddRADseq and Finstermeier-mtDNA datasets recovered *Callithrix* as the sistergroup of *Aotus*+*Saimiri*, while the UCE and exon-cap datasets recovered *Saimiri* as the sistergroup of *Aotus*+*Callithrix*; (2) the Finstermeier-mtDNA dataset placed *Papio* as sistergroup to *Cercocebus*+*Mandrillus*+*Macaca*, as opposed to *Cercocebus*+*Mandrillus* (other datasets). This topological uncertainty in the datasets is illustrated in Figure 4, with the ddRADseq dataset showing that the MCMC sampled a greater set of possible trees in this dataset; the UCE and exon-cap datasets had almost no topological discordance.

**Figure 4.**
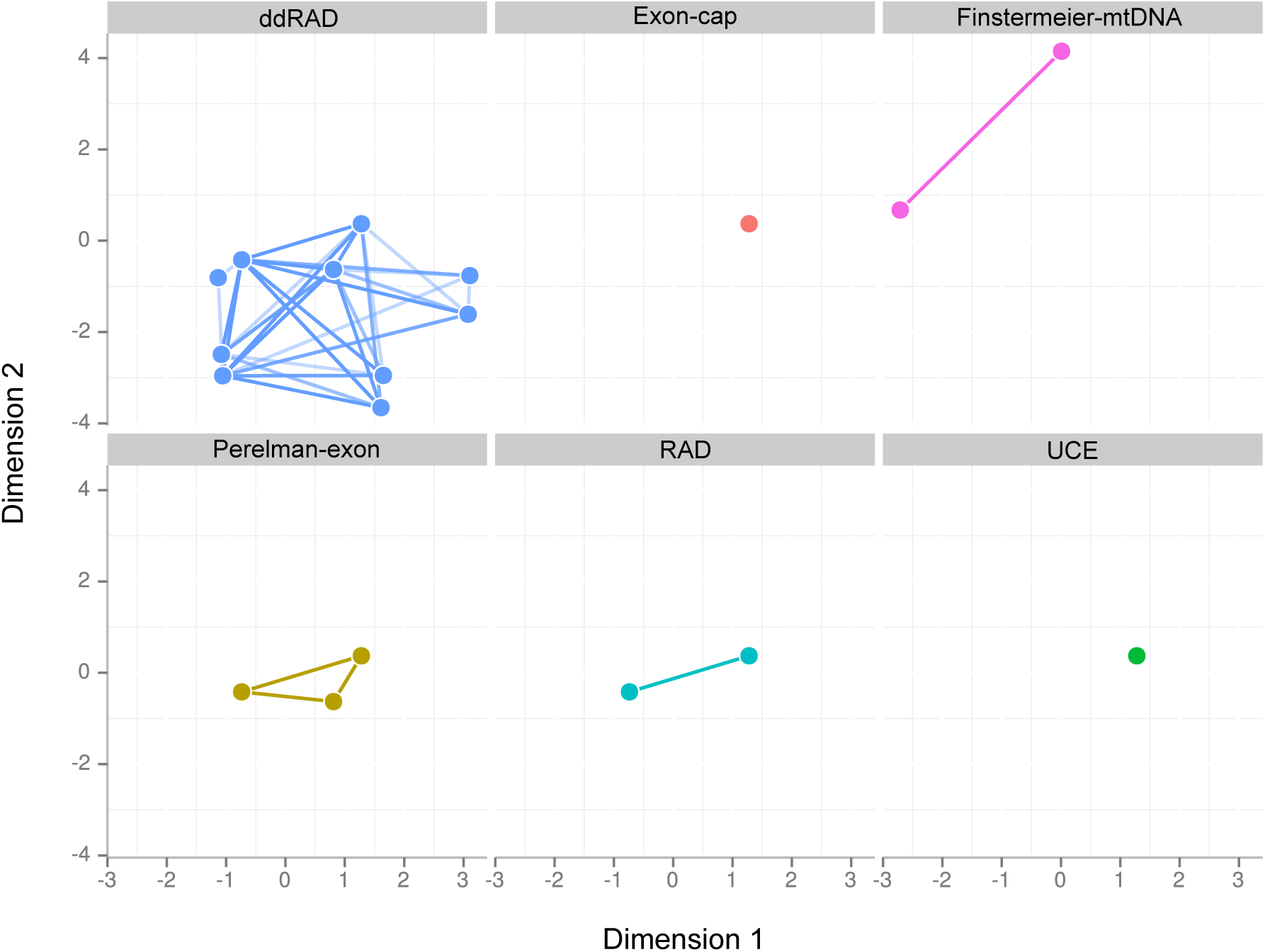
Phylogenetic uncertainty in a sample of 500 post-burnin ExaBayes trees for the six datasets. Treespace is shown using multidimensional scaling as implemented in the RWTY function *makeplot.treespace*. Only run one of two is shown, although the second run was checked for consistency with the first.

### Divergence times, parametric estimates and phylogenetic informativeness

A graphical summary of the dating analyses showing calibrated nodes, topology, and taxa is provided in Figure 1, while parameter estimates and statistical test results are presented in Table 5. Overall evolutionary rate estimates (substitutions per site, per Ma) varied from 0.17 (UCE) to 4.79 (Finstermeier-mtDNA). GC content estimates varied from 0.39 (UCE) to 0.51 (exon-cap). Among site rate variation as measured by the *α*-shape of the Γ parameter varied from 0.09 (exon-cap) to 0.57 (ddRADseq). The Φ*IC* criterion indicated greater support for the strict clock model over the relaxed clock model, with a Δ score greater than 39 for all datasets. The Finstermeier-mtDNA dataset was the least clocklike, with a likelihood ratio of 13.29 (Table 5).

**Table 5.**
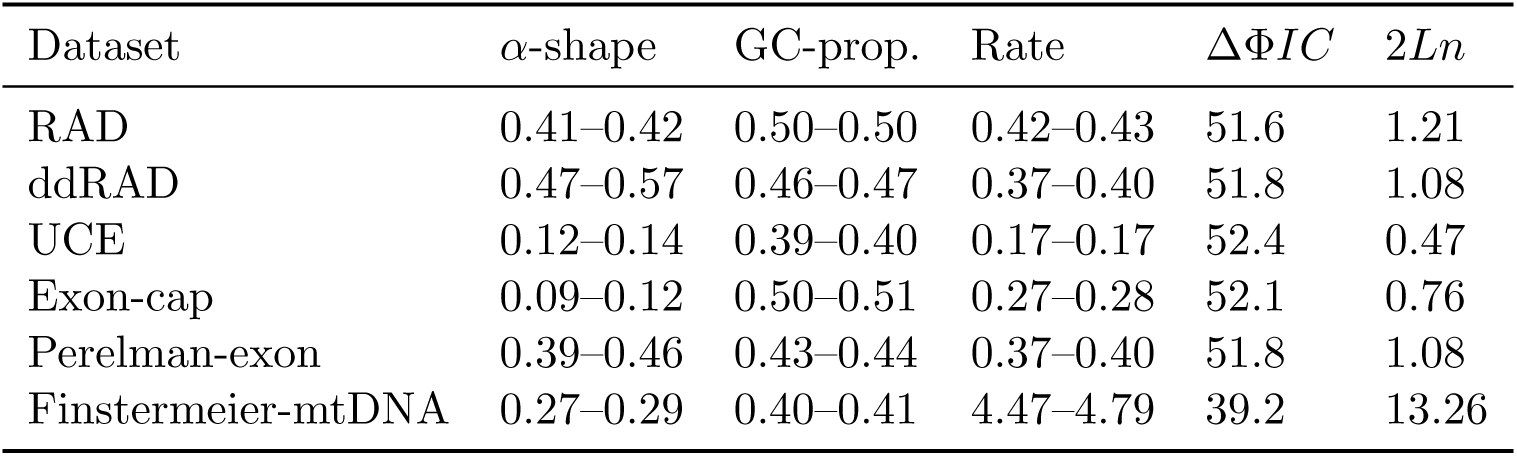
Highest posterior density (95% HPD) parameter estimates resulting from the ExaBayes analysis (9,000 post-burnin trees). Evolutionary rate as calculated from the sum of branch lengths from the posterior sample (number of substitutions per site per Ma). The clock-likeness of each dataset is estimated from the maximum clade credibility (MCC) tree using ΔΦ*IC* score of the relaxed clock model, and the likelihood ratio between the relaxed clock model and strict clock model (lower values indicate greater clocklikeness).

Divergence time estimates for selected clades under different clock models for each dataset is presented in Figure 5 (strict clock) and Figure 6 (relaxed clock), and shows that clade ages were estimated with considerably greater precision in the strict rather than the relaxed clock analyses. While ages estimated under the relaxed clock model were similar for all datasets, there were systematic differences among the strict clock dates, with the exon-cap and Perelman-exon datasets showing consistently younger ages than the UCE, RADseq, ddRADseq and Finstermeier-mtDNA for nodes < 30 Ma, but older and more uncertain age estimates, for the deepest nodes, e.g. Haplorrhini (Figure 5).

**Figure 5.**
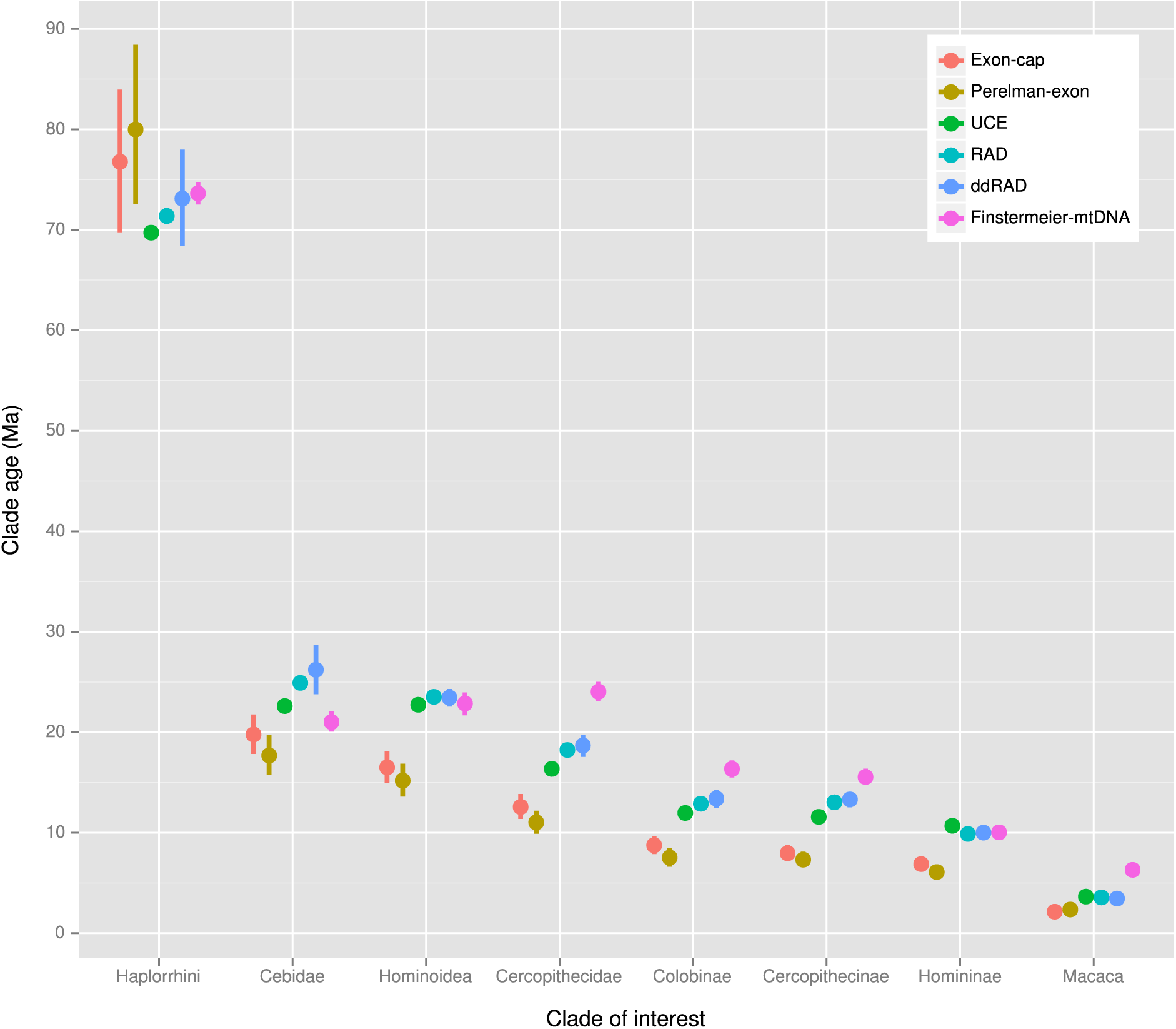
Divergence time estimates for selected clades from 9,000 post-burnin ExaBayes trees dated using the strict clock model implemented in the *chronos* function of Ape. Points show mean node heights; bars show 95% credible intervals.

**Figure 6.**
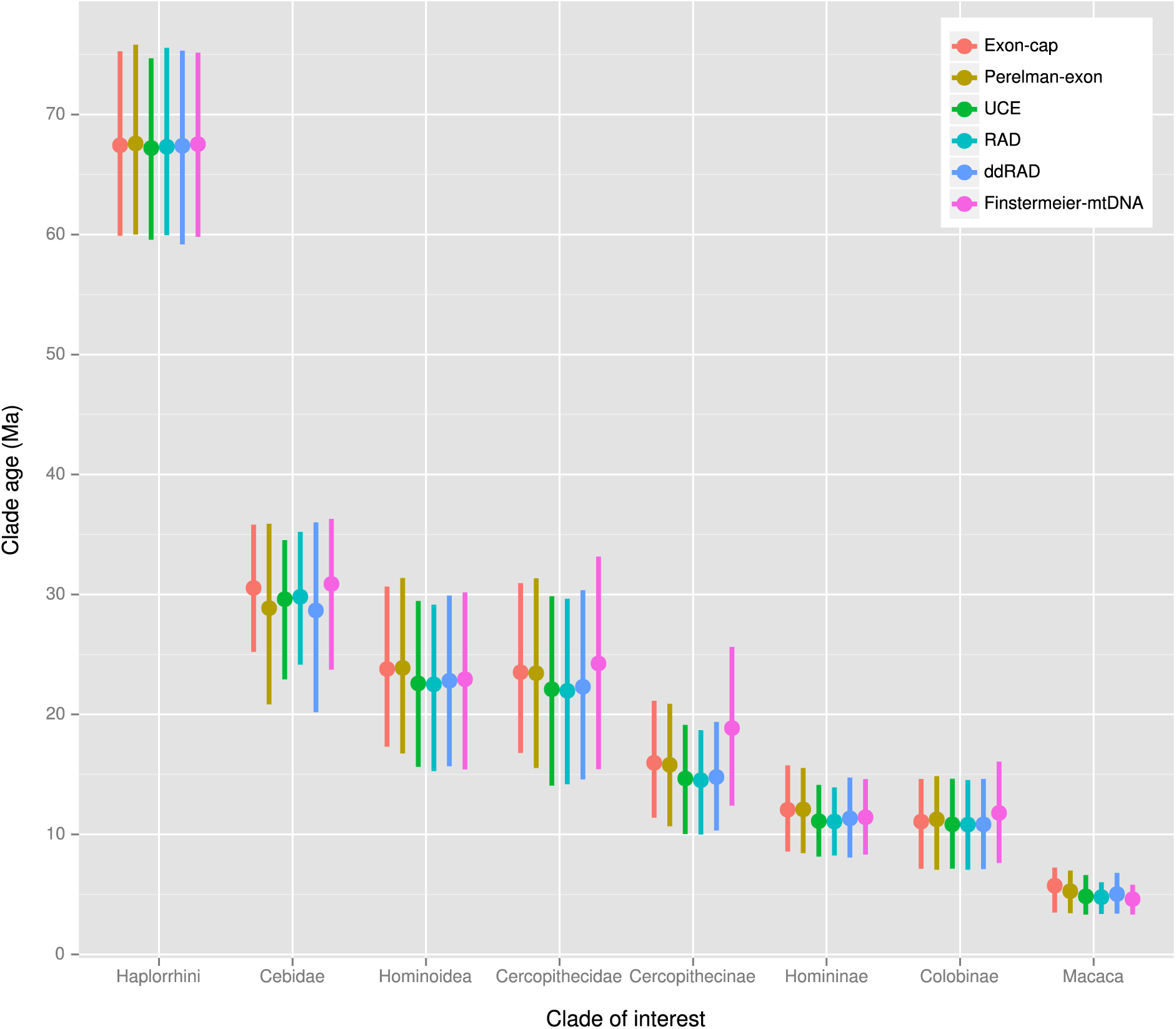
Divergence time estimates for selected clades from 9,000 post-burnin ExaBayes trees dated using the relaxed clock model implemented in the *chronos* function of Ape. Points show mean node heights; bars show 95% credible intervals.

Figure 7 shows phylogenetic informativeness (PI) over time. Peaks in PI can be seen at 5 Ma (Finstermeier-mtDNA), 21 Ma (ddRADseq), 22 Ma (RADseq), 32 Ma (UCE), 33 Ma (exon-cap), and 51 Ma (Perelman-exon). The RADseq, Finstermeier-mtDNA, ddRADseq and UCE datasets show more steeply sloping profiles than the exon-cap and Perelman-exon datasets, with these latter datasets not showing a rapid decay in PI either side of the peak. For comparison, log transformed and rescaled (per site) PI plots are shown in Supplementary Figure 8 and Supplementary Figure 7.

**Figure 7.**
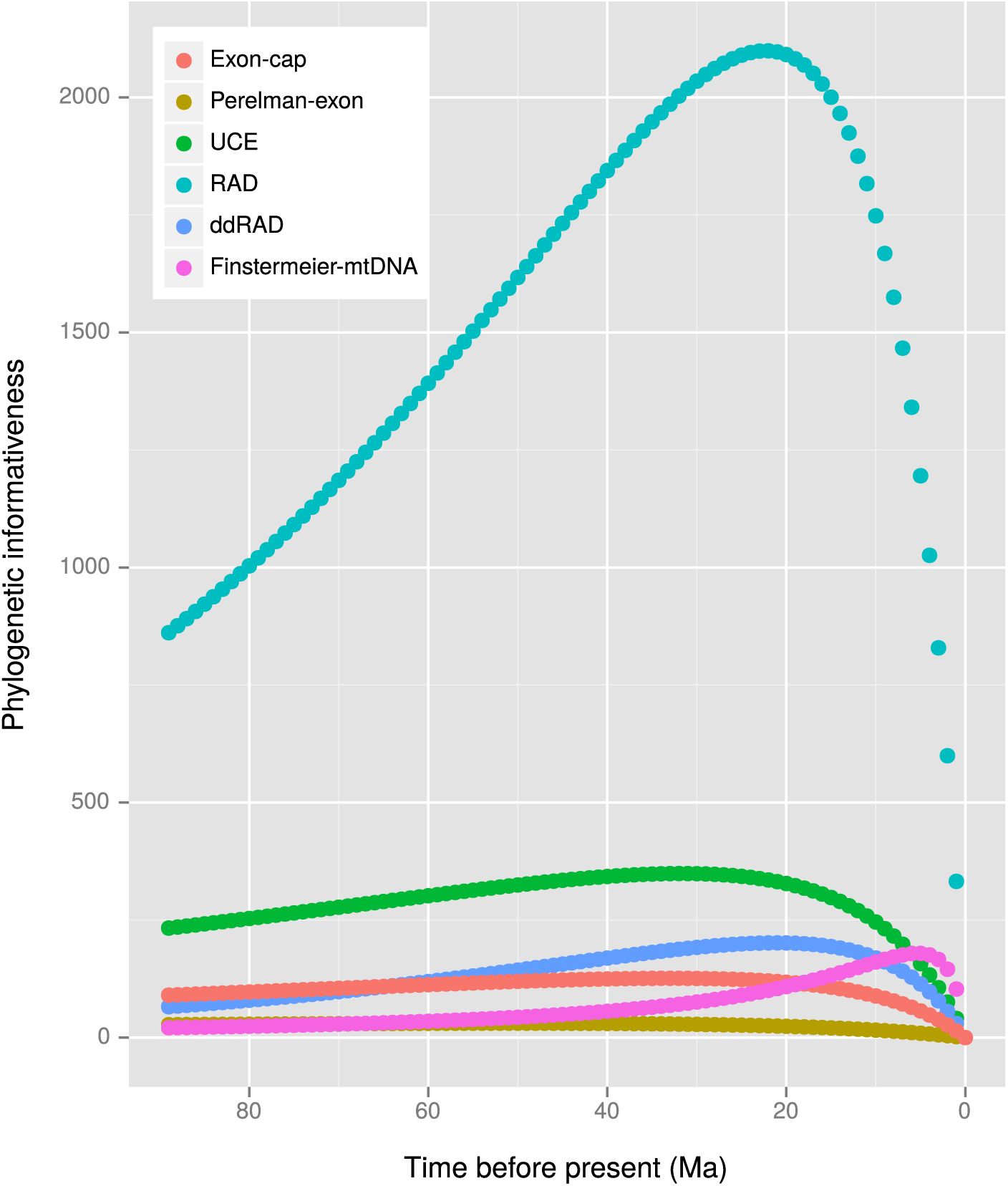
Phylogenetic informativeness (PI) over time for each dataset as calculated using Tapir. The strict clock MCC tree resulting from the analysis of the Perelman-exon dataset was used for the Tapir calculations.

## Discussion & summary

Our results show that the four protocols examined here are similar in their ability to estimate relationships and clade ages for a relatively tractable group. We examine below the main differences of the datasets in terms of phylogenetic/dating performance, and also in respect to analytical and data modelling issues, but we also provide an assessment and summary of the technical differences between the protocols in Appendix 1.

### Reduced genomic representation

Reduced representation protocols are universally applicable to non-model organisms since restriction sites are abundantly present in genomes of all organisms (Miller *et al*., 2007). The total number has a direct relationship with the size of the genome with approximately 6.9 recognition sequences per megabases reported for the 8-cutter *Sbfl* enzyme we used here (Herrera *et al*., 2015). Using the ddRADseq approach, one normally reduces the total number of RAD markers to one fifth to one tenth of the total, and therefore the total number of individuals that can be sequenced at the same coverage as in a RADseq experiment is roughly 10× to 20×. Although the dropout of ddRADseq loci paralleled RADseq, the total number of shared loci was lower (Figure 2; Figure 3), and this loss of loci came at the expense of phylogenetic resolution; our ddRADseq dataset was unable to resolve the basal relationships of the primates with confidence, and this was reflected in the uncertainty in the MCMC tree search (Figure 4). However, when the full ddRADseq dataset—6,202,215 sites, minimum four taxa per locus instead of 12—was analysed, the basal split received full support, although the cebid relationships continued to be not well supported (BPP 0.75 for *Saimiri*+*Aotus*). With the considerably larger RADseq dataset, all clades were recovered with unambiguous support. In terms of clade ages, while RADseq age distributions were marginally smaller than the ddRADseq dataset, both were consistent and overlapping (Figure 5, Figure 6). Both of these datasets also showed a similar pattern of relative phylogenetic informativeness (Supplementary Figure 8).

### Sequence capture

In theory the UCE and exon-cap protocols are universally or nearly universally applicable due to the conserved nature of the designed probes (Faircloth *et al*., 2012b; Li *et al*., 2013), but while probesets have been designed for phylogenetically broad groups (e.g. amniotes, vertebrates, teleost fishes), there can also be a substantial bioinformatic investment in probe design if one is interested in the less well-studied branches of the tree of life.

We found that both methods had a roughly similar proportions of recovered loci, at 45% (UCE) and 57% (exon-cap), after datasets were filtered to include no more than ≈ 50% missing taxa per locus. With filtering set at a minimum of four taxa per locus, more loci (2,771) were reported than probes (2,386) for the UCE dataset Table 2. This is due to the clustering step breaking up putatively orthologous sequences into more than one locus, with this being affected mainly as a result of the length of the incorporated flanking sequences; sequence variability increases with distance from the cores in both the UCE and exon-cap markers (Gilbert *et al*., 2015; Lemmon *et al*., 2012), so highly divergent intronic sites become difficult to cluster and align. Choosing shorter flanking sequences will result in a greater number of orthologous loci, but at the same time may lower per-locus phylogenetic informativeness. For the purpose of this study, we found a total length of 600 bp a reasonable compromise between number of loci and site variability.

In an *in vitro* experiment it is possible to control the length of the recovered loci by altering the sequencing library protocols to produce a larger insert size, up to, for example 1,700 bp (Lemmon *et al*., 2012). In practice, however, length of recovered loci is somewhat smaller. UCE studies using the standard tetrapod probe set achieved an average of 393 bp in birds and 412 bp in primates (Faircloth *et al*., 2012b), 604 bp in birds (Harvey *et al*., 2013), and 615 bp in lizards (Leaché *et al*., 2015). Using a different probeset for fishes, Faircloth *et al*. (2013) recovered an average UCE locus length of 457 bp. Harvey *et al*. (2013) also reported diminishing returns in species tree estimation at locus lengths greater than 500 bp. For exon-cap using the Lemmon *et al*. (2012) 512-locus probeset, Pyron *et al*. (2014) recovered a mean locus length of 676 bp in snakes, Leaché *et al*. (2014) a mean of 333 bp in lizards, and Eytan *et al*. (2015) a mean of 774 bp in fishes. Thus, our choice of 600 bp loci is typical of what can be expected from current technologies.

In terms of phylogenetic performance, both datasets performed identically in terms of phylogenetic performance, with all clades recovered with high support and no topological ambiguity (Figure 4). Although the total number of exon-cap loci was roughly one quarter of that of the UCE dataset, the exon-cap data had roughly double the proportion of parsimony informative sites (6% *versus* 3%; Table 3). Proportion of total missing sites was also lowest in the exon-cap dataset, as it showed the lowest locus dropout rate (Figure 2) and a more steady loss of loci as phylogenetic distance increased (linear *versus* exponential loss; Figure 3). Regarding clade ages, the exon-cap dataset consistently showed younger ages than the UCE data for recent nodes, but older ages for deeper nodes in the strict clock analysis. Reasons for these discrepancies are discussed below.

### Caveats & discrepancies

Our objective was not necessarily to assess accuracy—i.e. in recovering the true phylogeny or the true ages of primate clades—but rather to characterise whether the different datasets resulted in the same parameter estimates given identical analytical conditions. Therefore, we did not examine techniques that may improve the likelihood of recovering the true tree. Thus, we did not estimate species trees, we did not partition the data, we did not explore the effects of varying proportions of missing data, we used standard models, and we did not test the fit or adequacy of these models. Rather, we compared the consistency of our phylogenies to previously published results using independent datasets (Finstermeier *et al*., 2013; Perelman *et al*., 2011) and identical analytical conditions. We discuss briefly the justifications for doing so.

### Species trees and concatenated supermatrices

Despite the known theoretical advantages of models that account for gene tree heterogeneity over supermatrix concatenation methods (Edwards, 2009; Edwards *et al*., 2015; Kubatko & Degnan, 2007; McCormack *et al*., 2012a; Ogilvie *et al*., 2015), it has been suggested that these problems may not be universal to all datasets (see Roch & Warnow, 2015), and that supermatrix concatenation may display some advantage in making fewer assumptions about the data (Gatesy & Springer, 2014; Tonini *et al*., 2015).

Despite incomplete lineage sorting (ILS) being known in primates (Heckman *et al*., 2007; Mailund *et al*., 2014; Ting *et al*., 2008; Wang *et al*., 2012), overall our clades were well supported and almost entirely consistent between datasets. Therefore we did not suspect the stochastic effects of ILS to be having a pernicious effect on the results of our primate backbone phylogeny. However, three species—*Aotus, Callithrix*, and *Saimiri* representing the three subfamilies of the Cebidae, and which diverged from one another in rapid succession in the mid Miocene (Rosenberger, 2002)—were problematic and could not be resolved with satisfaction due to a short internodal branch at the base of family. Here, the conflicting but highly supported relationships (Supplementary Table 1) appear indicative of a mismodelling of gene tree heterogeneity. Testing which relationship is ultimately correct—i.e. RADseq, ddRADseq, Finstermeier-mtDNA *versus* UCE, exon-cap, Perelman-exon, non-genic markers (Jameson Kiesling *et al*., 2014)—will require greater taxon sampling and biologically more appropriate models incorporating the multispecies coalescent (Heled & Drummond, 2010). This will allow us to gauge which of the datasets is being mismodelled, and whether the true signal can be extracted under the correct analytical conditions. A third well-supported topology (*Aotus* sistergroup to *Callithrix*+*Saimiri*) has also been reported (Supplementary Table 1; Wildman *et al*., 2009).

We additionally highlight that the mitochondrial genome dataset (Finstermeier-mtDNA) was inconsistent with the nuclear genomic consensus in respect to the position of *Papio anubis*, and therefore we suggest caution in using single locus datasets, given that gene trees may not represent the species tree (Maddison, 1997).

### Data partitioning and branch length estimation

Partitioning of data into subsets of similarly evolving sites is another frequently used method for addressing heterogeneity in empirical data (Brandley *et al*., 2005; Brown & Lemmon, 2007). It was reported by Kainer & Lanfear (2015) that suboptimal partitioning mostly affected topologies at uncertain nodes (e.g. the Cebidae), but also that branch lengths, and therefore clade ages, could be poorly estimated if the models were not sufficiently complex. However, these effects appeared to decrease as datasets got larger, and all of our datasets are considerably greater in size than those tested by Kainer & Lanfear. In fact, due to the large quantitity of data, branch lengths are often estimated in genomic datasets with very small error (Table 5; Ho, 2014), and age estimates will depend more on the clock model and even more on the number and quality of fossil calibrations (Warnock *et al*., 2014; Zheng & Wiens, 2015).

Our results show that the dates estimated were sensitive to clock model used, with the strict clock having considerably smaller credible intervals, but estimating generally younger ages than the relaxed clock for the more recent nodes (< 30 Ma). Despite the findings of Lepage *et al*. (2007), where empirical data consistently departed from the strict clock in favor of autocorrelated or uncorrelated models, we found that none of our datasets could reject the null model of a single rate. However, rate variation in primates is well documented (Heath *et al*., 2012; Jameson Kiesling *et al*., 2014; Peng *et al*., 2009), and more sophisticated Bayesian methods (e.g. Drummond & Suchard, 2010; Heath *et al*., 2012) may be able to discern whether this is a general pattern of genome scale datasets or an analytical artefact. Indeed, Paradis (2013) reported that the Φ*IC* appears a conservative test, with it sometimes rejecting autocorrelated and relaxed clocks when the data were simulated under those models.

Discrepancies between the data classes were also apparent, with the exon datasets (exon-cap, Perelman-exon) estimating consistently younger ages than the RADseq, ddRADseq, UCE and Finstermeier-mtDNA datasets. As demonstrated by Dornburg *et al*. (2014), we suggest that this discrepancy can also be explained by patterns in the phylogenetic informativeness of the markers, with the datasets showing PI profiles strongly peaking toward the present, tending to estimate older dates for the more recent nodes and younger dates for the older nodes (Figure 5, Figure 7). This corresponds to “tree extension/compression” errors in branch-length estimates *sensu* Phillips (2009), whereby hidden substitutions older than the peak PI are increasingly underestimated due to sequence saturation, and hidden substitutions close to the age of the peak PI are overestimated. This phenomenon can be exasperated when combined with the strict clock model, which fits a constant-rate linear-model to time heterogeneous data. Therefore, phylogeneticists should be aware of the dangers of false precision in molecular dating (Graur & Martin, 2004), as well as the influences of clock models (Paradis, 2013), fossil calibrations (Parham *et al*., 2012; Pozzi *et al*., 2014; Warnock *et al*., 2014), and patterns in molecular evolution (Dornburg *et al*., 2014; Ho & Duchene, 2014; Ho, 2014; Phillips, 2009).

### Missing data

Due to mutations at restriction sites the dropout rate of traditionally-generated RADseq loci increases dramatically with phylogenetic divergence (Figure 2; Figure 3; Cariou *et al*., 2013), causing a highly uneven distribution of missing data. Wagner *et al*. (2013) demonstrated that if the permitted proportion of missing data is set too low, support and resolution can be eroded because otherwise informative and variable loci are discarded (Huang & Knowles, 2014; Arnold *et al*., 2013). But conversely, as increasing amounts of missing data are included the benefits can diminish (Streicher *et al*., 2015). The effects at deep phylogenetic levels can be seen in the ddRADseq analysis, which when set at ≈ 50% missing data, very few loci were available to resolve the basal nodes (Table 4, Figure 4, Supplementary Figure 5). When set at a maximum of ≈ 80% missing taxa per locus, the support increased for the uncontroversial basal relationships, although it must be noted that support for the controversial cebid node did not improve.

Although we standardized the analysis by fixing the level of missing taxa *a priori* to a seemingly arbitrary proportion of ≈ 50% at any given locus, this value was reported by Streicher *et al*. (2015) as showing the maximal level of support in their analyses, and it appears to be a good compromise value. However, the effect of missing data may be ultimately be unequal between our datasets due to the non-random nature of this missing data (Hovmöller *et al*., 2013; Shavit Grievink *et al*., 2013; Roure *et al*., 2013). Therefore, it is important to note the difference between the amount of missing taxa per locus, and the overall proportion of missing sites (Streicher *et al*., 2015); all of our datasets had the same number of missing taxa per locus, but varied considerably in the overall proportion of missing data (Table 3), due to the unevenness of where the loci are dropping out (Figure 2). The UCE and exon-cap datasets were considerably less affected by uneven distribution of loci than the RADseq datasets.

### Bioinformatics and other issues

An additional, but important aspect that we did not consider fully is that of the settings of the bioinformatic assembly parameters. In particular, the clustering step that establishes sequence orthology is known to significantly influence the total number and taxon composition of recovered RAD loci (Cariou *et al*., 2013; Leaché *et al*., 2015; Rubin *et al*., 2012). Applying more liberal conditions, i.e. a lower sequence similarity threshold, will result in fewer loci and consequently a greater number of shared loci, but can introduce what Leaché *et al*. (2015) term “RAD noise”, stemming from increasing frequency of treating paralogous loci as orthologous (Cariou *et al*., 2013). This noise led to reported inconsistencies between ddRADseq and UCE results of Leaché *et al*. (2015), with clades associated with short internal branches being inconsistently recovered by the ddRADseq datasets. However, Leaché *et al*. used ddRAD data comprising 39 bp sequences *versus* UCE data with mean contig length of 615 bp, and the RAD noise reported is more a consequence of the difficulty of establishing orthology of short-read sequences (Cariou *et al*., 2013), than a fundamental property of RAD sequence data. Therefore when using “traditionally-generated” RADseq loci—i.e. 50–100 bp Illumina reads before removing barcode and restriction site sequences—establishing orthology for such short sequences will be a much greater problem that for longer RAD sequences generated from paired-end Illumina sequencing experiments, or the 300–400 bp reads generated on the IonTorrent PGM.

Despite this, Rubin *et al*. (2012) reported benefits from the greater number of loci recovered with more liberal settings, and showed that even when clusters are not fully orthologous they will nevertheless contain phylogenetic information. Therefore it appears that the influence of this parameter is dependant on tree shape and phylogenetic informativeness of the loci, so for ease of comparison we left the similarity threshold at an intermediate value (PyRAD default of 88%), but for empirical studies, and particularly for trees with short internodes, we encourage a more thorough examination of the effects of changing these settings (see Leaché *et al*., 2015).

As this was an *in silico* study, we were also unable to control for some aspects of an *in vitro* lab experiment, and in particular we did not explicitly model whether our MegaBlast “capture” of the UCEs and exons reflected hybridisation conditions *in vitro*. However, the proportion of recovered loci was roughly consistent with empirical studies: 333 loci (Pyron *et al*., 2014, exon-cap), 215 (Leaché *et al*., 2014, exon-cap); 107 (Eytan *et al*., 2015, exon-cap); 1145 (Crawford *et al*., 2012, UCE); 854 (Faircloth *et al*., 2012b, UCE); and 776–1516 (Smith *et al*., 2014, UCE). We also did not consider variable template quality and its effect on enrichment success and sequencing depth (McCormack *et al*., 2015).

It is also important to note that we only examined published protocols, and so results may not necessarily be completely representative of that general type of nucleotide data. New methods are continuing to be developed, and new probesets for additional taxonomic groups have been published (http://ultraconserved.org/, http://anchoredphylogeny.com/). Alternatively, custom probes can also be designed (Bragg *et al*., 2015).

## Conclusions & recommendations

In spite of these caveats, the phylogenetic hypotheses based on the four different genomic datasets were largely non-conflicting with each other, and also with respect to the independent multi-gene and mitogenomic datasets of Perelman *et al*. (2011) and Finstermeier *et al*. (2013), respectively. While the modest number of ddRADseq loci sampled in early diverging species had insufficient power to resolve deeper nodes, all marker types examined show the same ability to robustly reconstruct phylogenetic relationships of primates, both recent as well as ancient (up to ≈ 80 Ma). The temporal breadth of the primate phylogeny and the diversity of ancient as well as recent speciation events probably will represent the range of phylogenetic questions addressed by a great majority of systematists and evolutionary biologists. Therefore, our conclusions should not be viewed as being restricted to primates, but can be viewed as general and most likely broadly applicable to many taxonomic groups.

However, genome scale data do not magically resolve *all* phylogenetic relationships, and cannot be considered a panacea (Philippe *et al*., 2011; Pyron, 2015). For certain classic, phylogenetically hard-to-resolve problems such as ancient, rapid radiations or extreme rate variation, only much improved models and data quality assessments may be able to reduce incongruence or avoid systematic biases in these cases (Doyle *et al*., 2015; Jarvis *et al*., 2014; Jeffroy *et al*., 2006; Phillips *et al*., 2004; Rodríguez-Ezpeleta *et al*., 2007). Here we show an example of this in the case of the Cebidae, whereby completing hypotheses were recovered with high support by different datasets. It is therefore essential that any type of genomic data be explored thoroughly and tested for the effects of competing assumptions before strong conclusions are made (even if particular phylogenetic hypotheses are strongly supported). This is especially true regarding the estimation of clade ages with genomic data. Although the uncorrelated relaxed-clock model estimated divergence times that were almost identical between the datasets, the strict clock model reported extremely precise but inconsistent results and further investigation into the appropriateness of this model and the factors influencing it is required.

When compared to the results from the sequence capture methods, it is possible that the RADseq protocol generated more data than was actually necessary for resolving the phylogeny over the time scales studied here. However, the RADseq and ddRADseq data also have much higher relative, and in the case of RADseq also absolute information content, and thus are likely a better choice for resolving relationships at the population to species boundaries. Furthermore, in instances where investigators only have access to a low capacity NGS machine such as the IonTorrent PGM, the ddRADseq protocol offers best value for money since an entire experiment can be carried out in one or two sequencing runs without compromising data quality or phylogenetic resolution at the time scales investigated here.

Comparing the UCE and exon-cap protocols, the latter provided the most complete data matrix, was least affected by phylogenetic divergence between taxa, and also displayed the most reliable, constant rate PI profile for molecular dating. With their greater degree of standardization and lower anonymity of the loci, both protocols also offer a more reliable solution to data sharing. For datasets where a large number of individuals need to be sequenced, the 512-locus exon-cap probe-set represents one fifth of the potential number of loci of the UCE protocol, and given that phylogenetic performance was similar with these fewer loci, and the exon-cap data retains its phylogenetic informativeness at deeper phylogenetic divergences, it would be possible to expand taxon sampling considerably. Unfortunately, enthusiasm for coding regions as phylogenetic markers may be tempered by the results of Jarvis *et al*. (2014), who reported consistent incongruence between their favored trees and their exon-only trees, and particularly so for a high variance subset of coding sites comprising non-stationary third positions. Although this issue appears more relevant to the hardest-to-resolve parts of the tree, it reinforces the need to understand potential sources of systematic error in each dataset (Eytan *et al*., 2015; Philippe *et al*., 2011; Rodríguez-Ezpeleta *et al*., 2007).

In summary, any of the data types analyzed in this study will confidently resolve phylogenetic relationships over the evolutionary time scales studied here due to the preponderance of massive amounts of phylogenetically informative data. Thus ultimately the choice of phylogenomic protocol should depend on a laboratory’s existing expertise, equipment and funding, phylogenetic breath of a laboratory’s projects, as well as the necessity to generate standardized datasets in collaborative projects, or for comparison with existing data.

## Acknowledgements

We thank CNPq for “Science without Borders” post-doctoral fellowship to RAC and a senior post-doctoral fellowship to TH. The CNPq/SISBIOTA-BioPHAM (grant no. CNPq 563348/2010) and the NSF/FAPESP “Dimensions of Biodiversity” (grant nos. NSF 1241066 and FAPESP 12/50260–6) projects provided additional support.

---

**Supplementary Figure 1.**
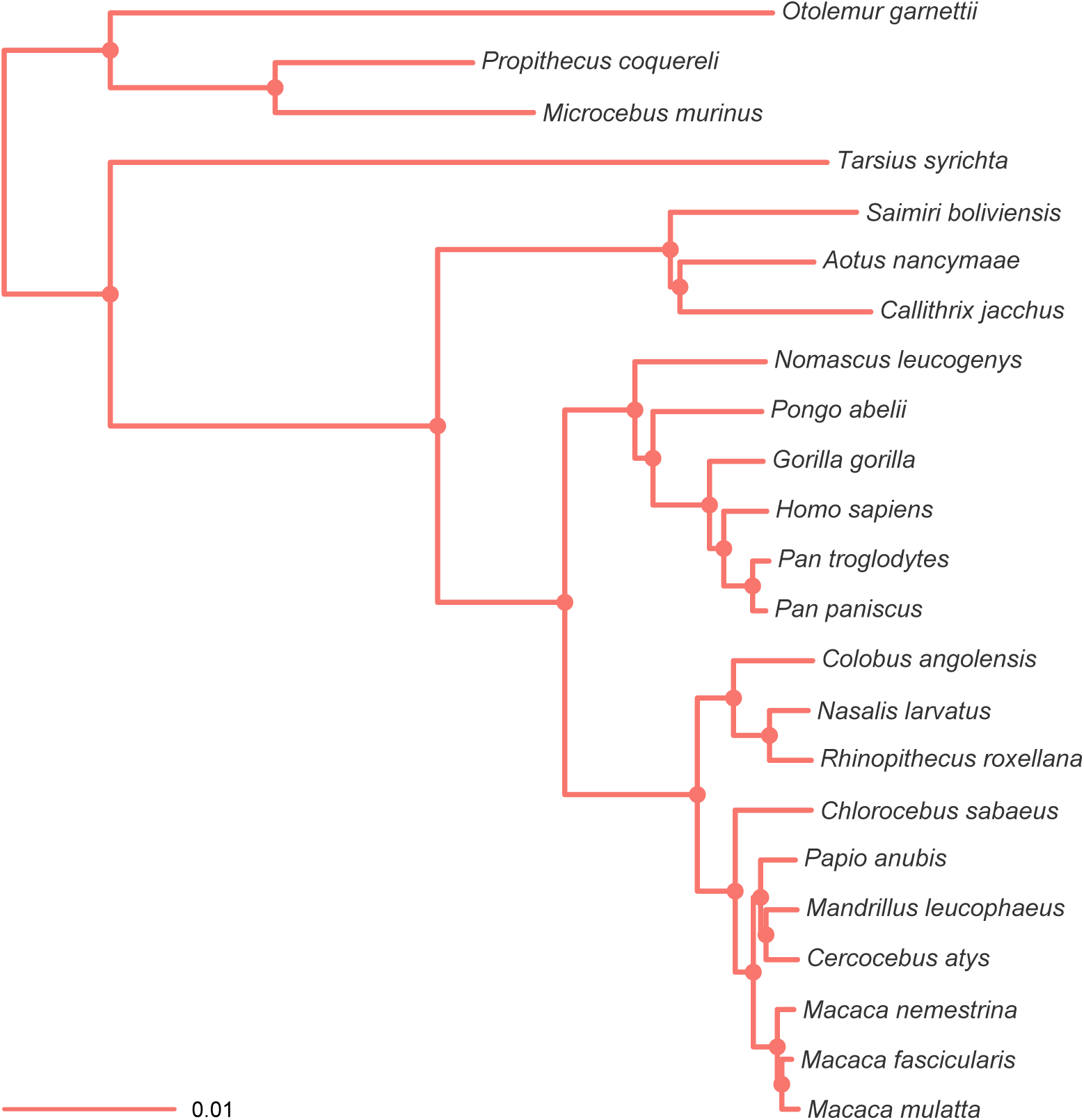
Maximum clade credibility (MCC) tree resulting from the ExaBayes analysis of the exon-cap dataset. Tree was rooted on the branch leading to *Otolemur, Propithecus* and *Microcebus*. Nodes are colored according to Bayesian posterior probability (BPP), with filled nodes having BPP ≥ 0.99; open nodes are labelled with BPP value.

**Supplementary Figure 2.**
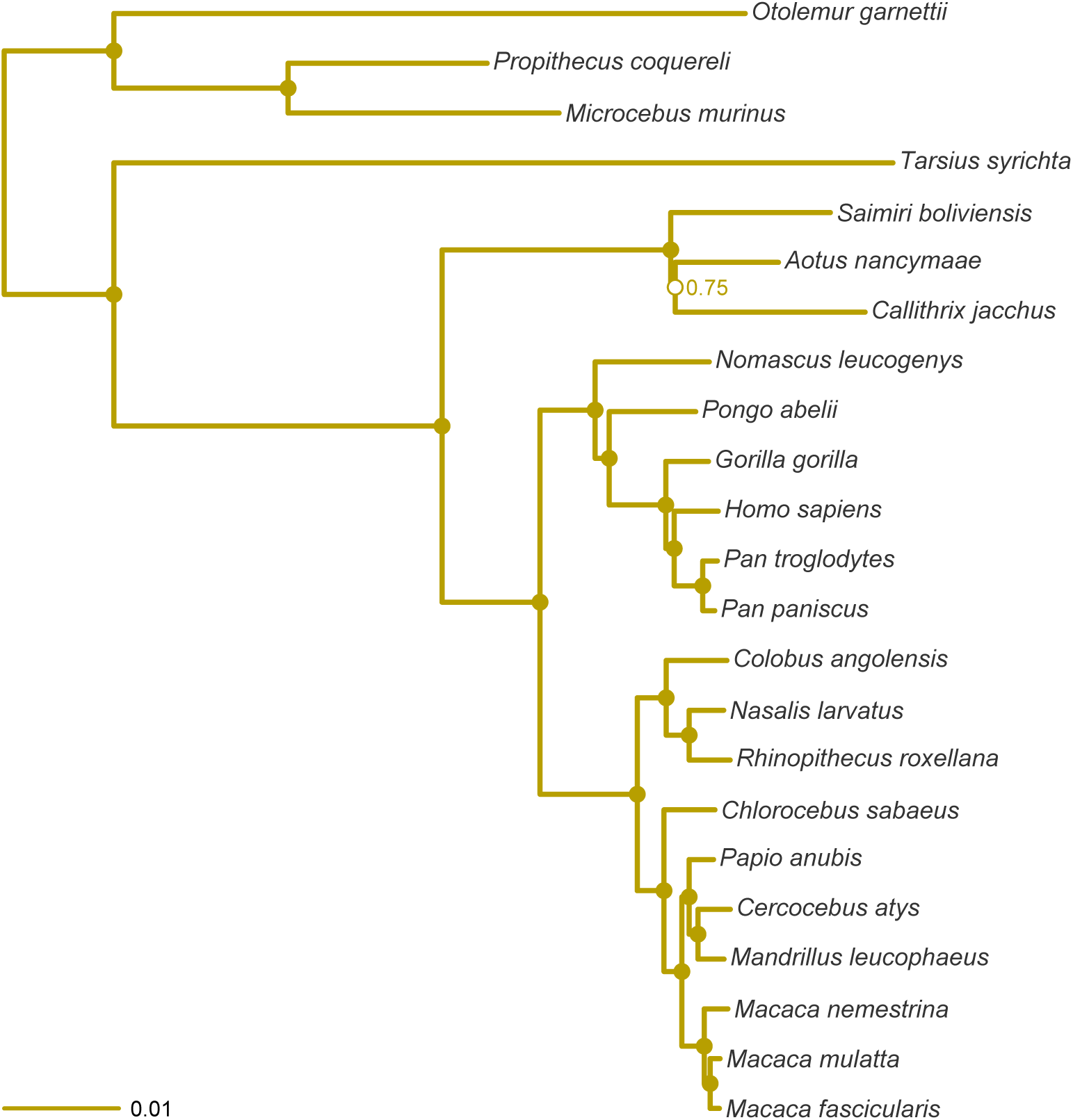
MCC tree resulting from the ExaBayes analysis of the pruned Perelman-exon dataset. Tree was rooted on the branch leading to *Otolemur, Propithecus* and *Microcebus*. Nodes are colored according to Bayesian posterior probability (BPP), with filled nodes having BPP ≥ 0.99; open nodes are labelled with BPP value.

**Supplementary Figure 3.**
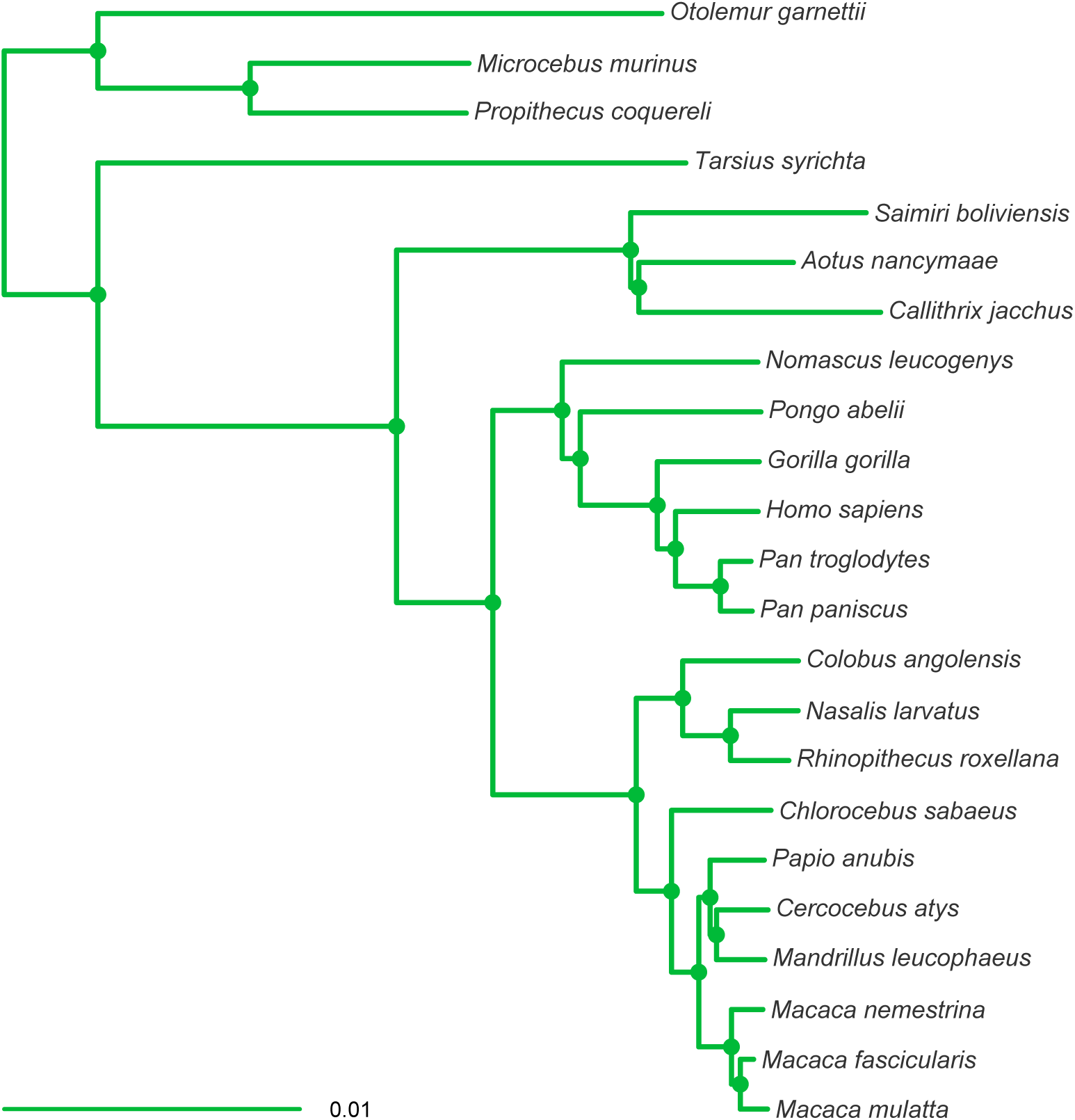
MCC tree resulting from the ExaBayes analysis of the UCE dataset. Tree was rooted on the branch leading to *Otolemur*, *Propithecus* and *Microcebus*. Nodes are colored according to Bayesian posterior probability (BPP), with filled nodes having BPP ≥ 0.99; open nodes are labelled with BPP value.

**Supplementary Figure 4.**
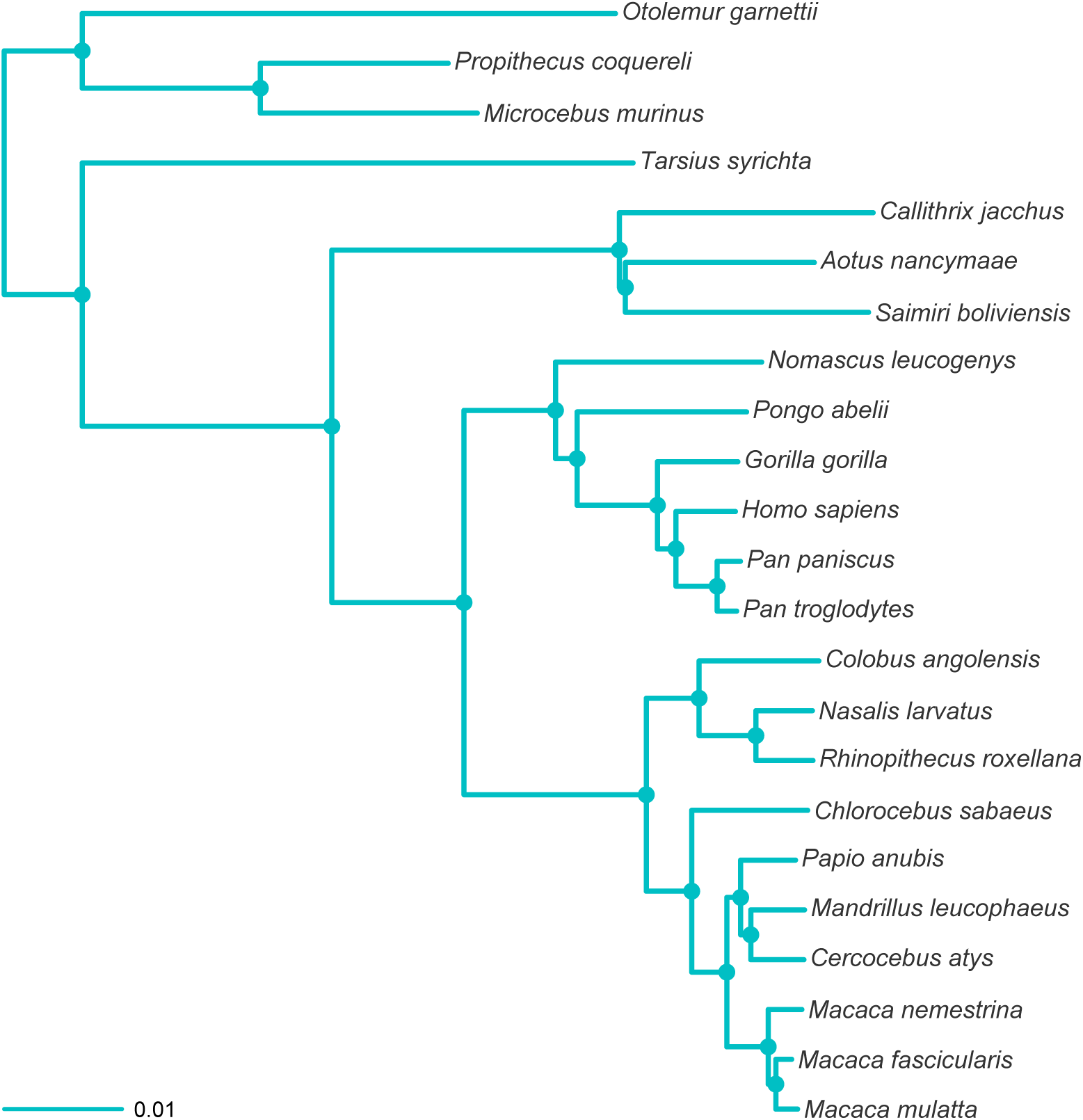
MCC tree resulting from the ExaBayes analysis of the RADseq dataset. Tree was rooted on the branch leading to *Otolemur, Propithecus* and *Microcebus*. Nodes are colored according to Bayesian posterior probability (BPP), with filled nodes having BPP ≥ 0.99; open nodes are labelled with BPP value.

**Supplementary Figure 5.**
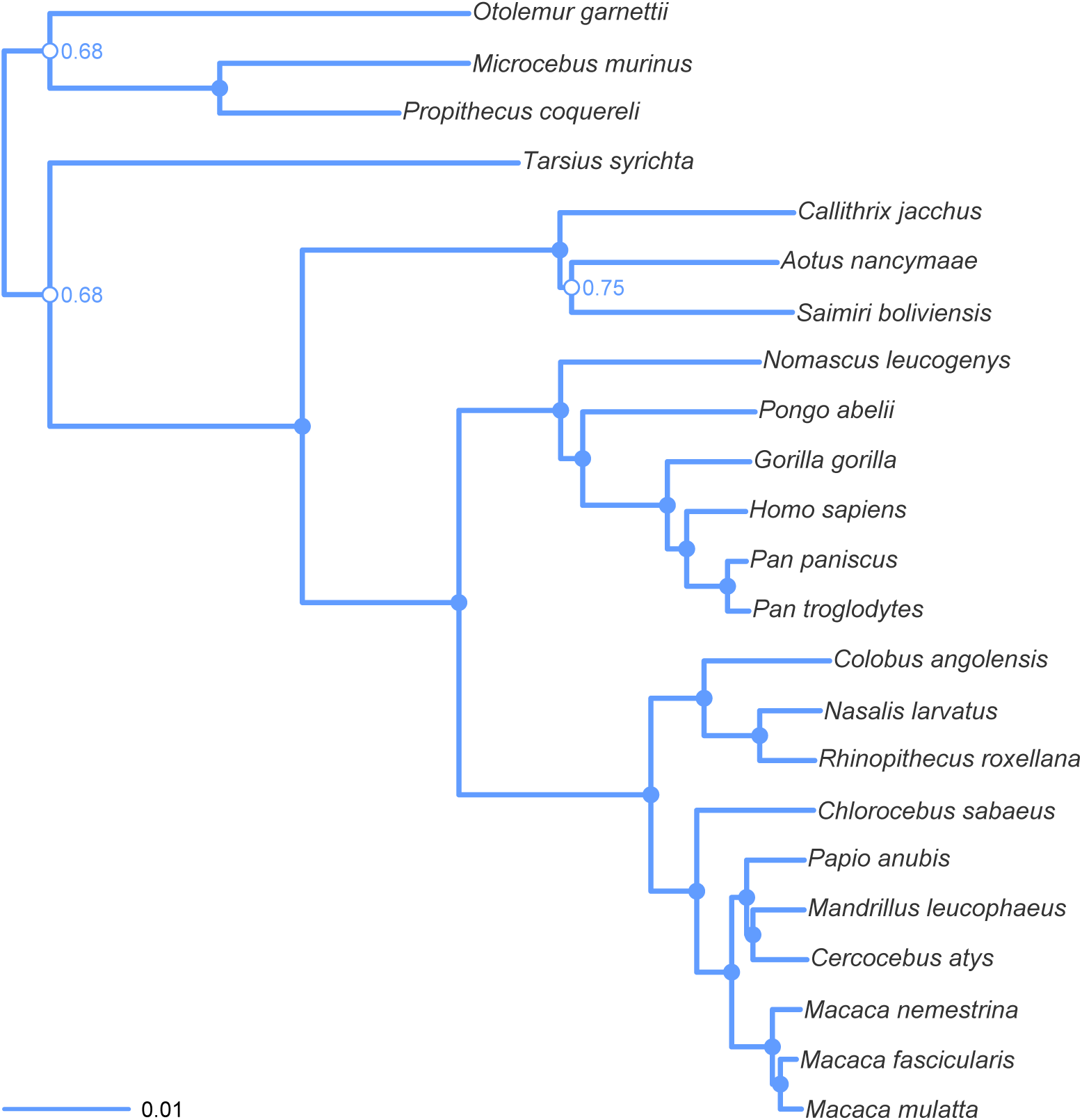
MCC tree resulting from the ExaBayes analysis of the ddRADseq dataset. Tree was rooted on the branch leading to *Otolemur, Propithecus* and *Microcebus*. Nodes are colored according to Bayesian posterior probability (BPP), with filled nodes having BPP ≥ 0.99; open nodes are labelled with BPP value.

**Supplementary Figure 6.**
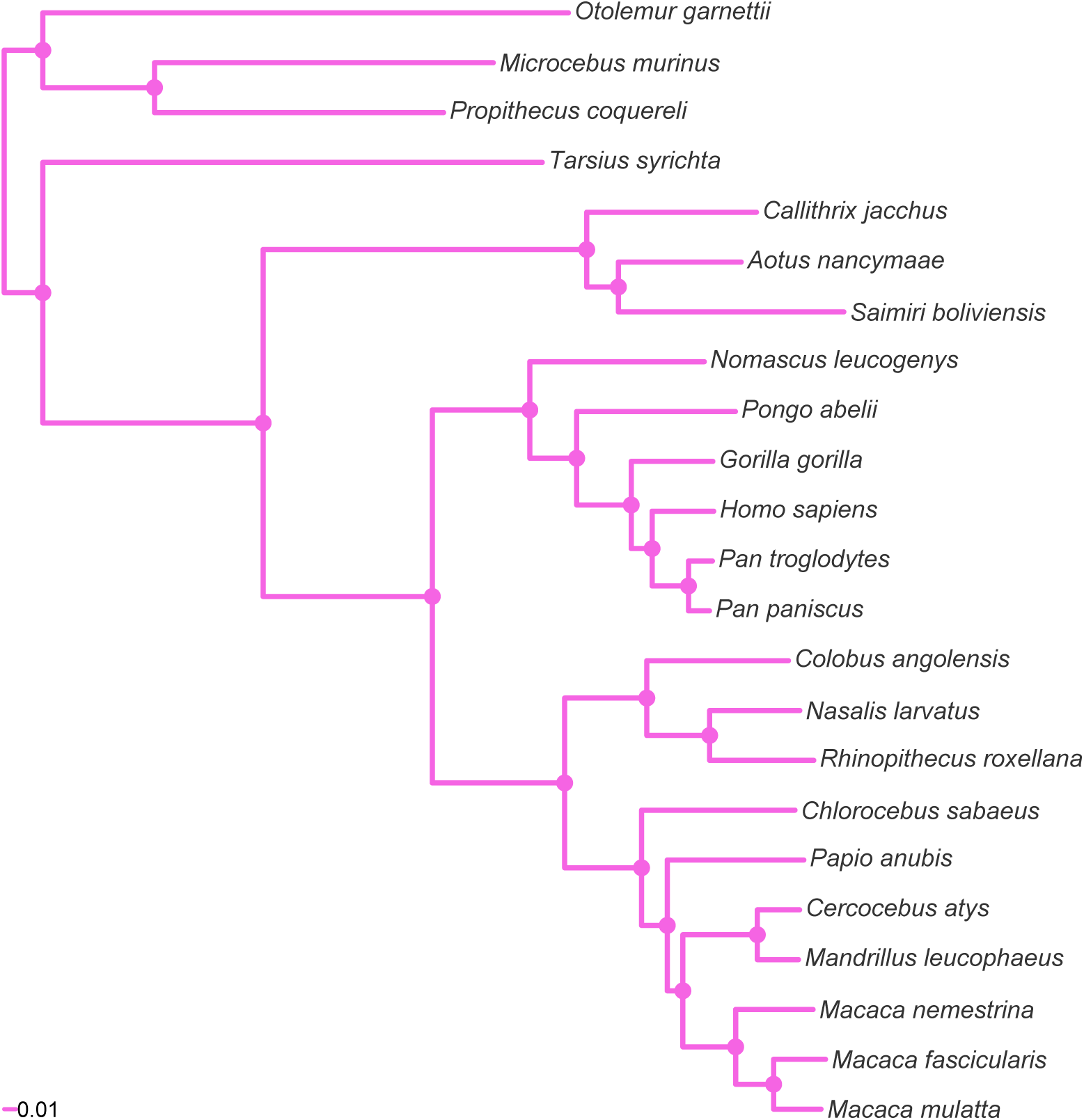
MCC tree resulting from the ExaBayes analysis of the pruned Finstermeier-mtDNA dataset. Tree was rooted on the branch leading to *Otolemur, Propithecus* and *Microcebus*. Nodes are colored according to Bayesian posterior probability (BPP), with filled nodes having BPP ≥ 0.99; open nodes are labelled with BPP value.

**Supplementary Figure 7.**
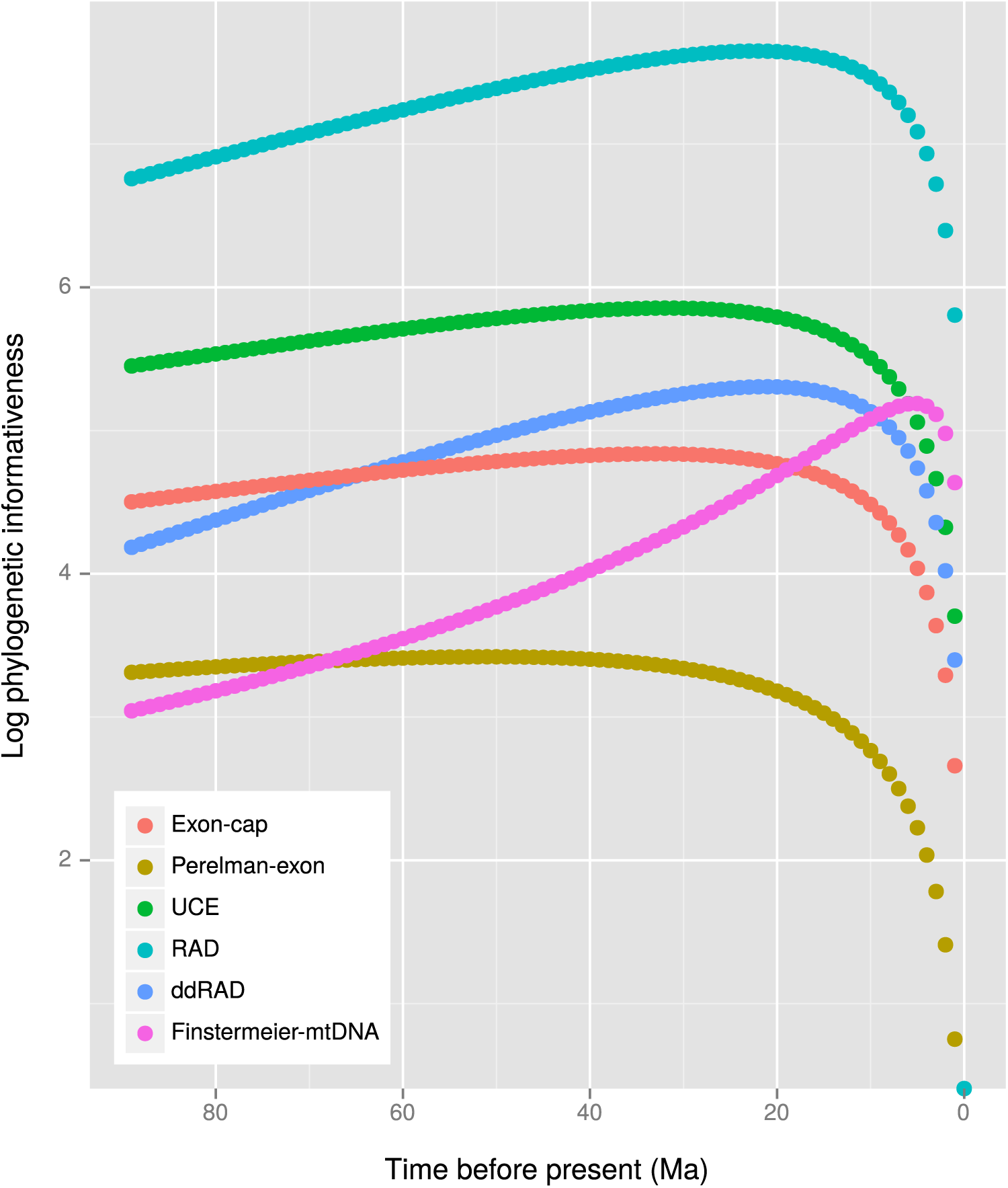
Log phylogenetic informativeness (PI) over time for each dataset as calculated using Tapir; The strict clock MCC tree resulting from the analysis of the Perelman-exon dataset was used for the Tapir calculations.

**Supplementary Figure 8.**
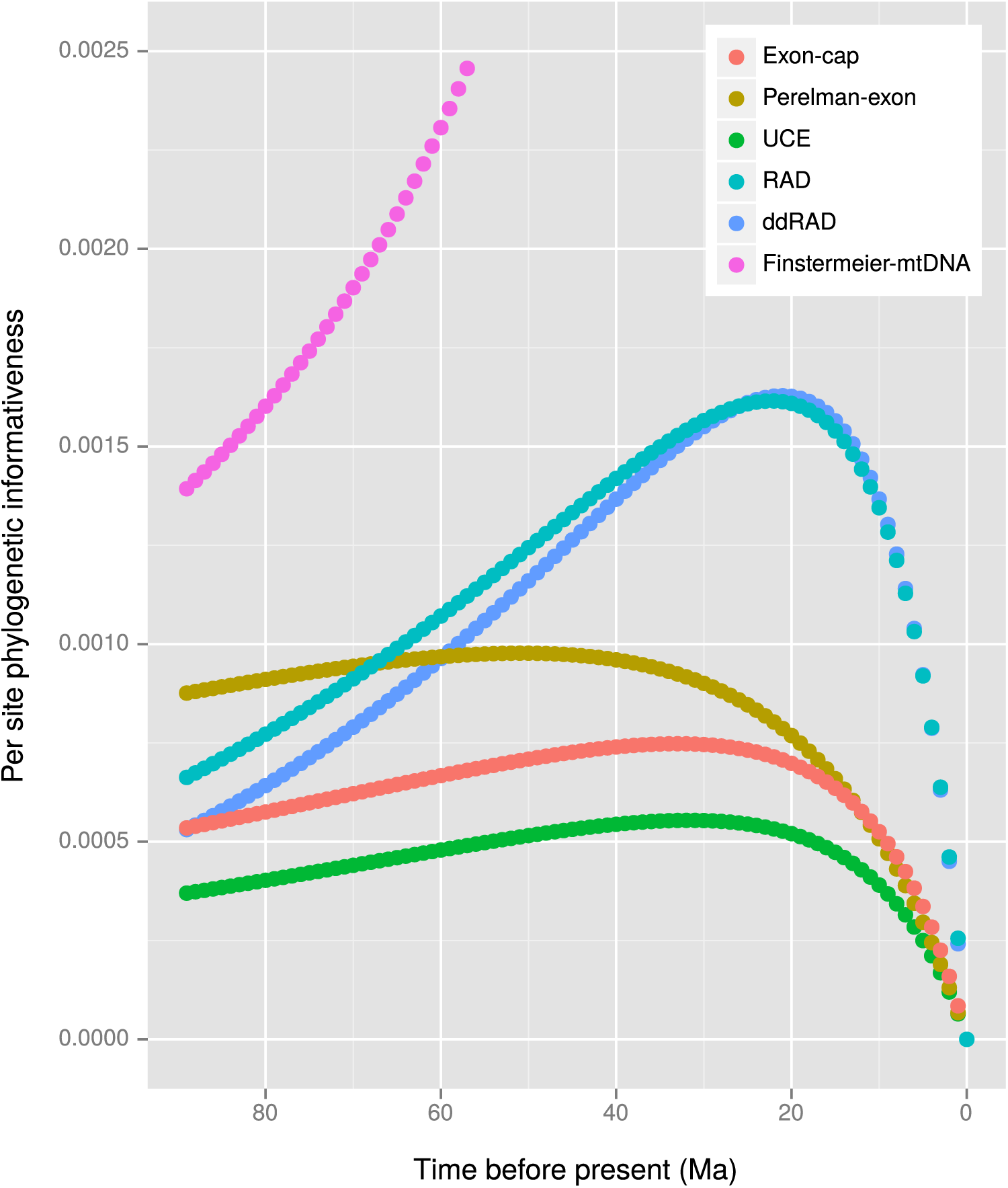
Relative PI over time for each dataset as calculated using Tapir; PI was scaled to the total number of sites in each dataset (Table 3) to enable better comparison of information content. Note that the PI goes off the scale for the Finstermeier-mtDNA dataset, peaking at PI of 0.012 at approximately 5 Ma. The strict clock MCC tree resulting from the analysis of the Perelman-exon dataset was used for the Tapir calculations.

**Supplementary Table 1.** Summary of support (Bayesian posterior probability) from the different datasets/studies for alternate phylogenetic relationships among the Cebidae. Species names are abbreviated as follows: *Aotus* (*Ao*), *Callithrix* (*Ca*), and *Saimiri* (*Sa*).

| Dataset | *Ca*,(*Ao*,*Sa*) | *Sa*,(*Ao*,*Ca*) | *Ao*,(*Sa*,*Ca*) |
| --- | --- | --- | --- |
| RAD | 0.99 | - | - |
| ddRAD | 0.75 | - | - |
| UCE | - | 1.00 | - |
| Exon-cap | - | 1.00 | - |
| Perelman-exon (this study) | - | 0.75 | - |
| Perelman *et al.* (2011) | - | 1.00 | - |
| Finstermeier-mtDNA (this study) | 0.99 | - | - |
| Finstermeier *et al.* (2013) | - | 0.72 | - |
| Jameson Kiesling *et al.* (2014) | - | 1.00 | - |
| Wildman *et al.* (2009) | - | - | 0.99 |

## Appendix 1: technical comparison

All protocols rely on basic laboratory equipment such as PCR machines and gel electrophoresis equipment. In the RAD, UCE and exon-cap protocols one needs to shear DNA which requires either a sonicator (e.g. Bioruptor) or a nebulizer (e.g. Covaris). The enzyme fragmentase can also be used for DNA shearing. In all protocols, there a need to size select the DNA fragments for sequencing, however, size selection is especially critical in the ddRADseq protocol if one wants to combine different runs, and therefore PippinPrep or BluPippin is recommended.

All protocols also require an *a priori* investment into reagents to prepare the library. For the RADseq and ddRADseq protocol this represents restriction and modifying enzymes, while for UCE and exon-cap protocols this represents investment in hybridization probes and modifying enzymes. All protocols will also require the purchase of barcoded and phosphorylated adaptors and PCR primers. The purchase of phosphorylated adaptors and PCR primers can run into several thousand dollars depending on the barcoding strategy used, while restriction and modifying enzymes will cost several hundred dollars. The quantity of adaptors, primers and enzymes are, however, sufficient for the preparation of several thousands of samples. UCE hybridization probes need to be purchased for each samples to be enriched, and cost between $75 and $30 per sample depending on the quantity of probes purchased (see http://www.mycroarray.com/mybaits/mybaits-UCEs.html). Exon-cap probes will need to be custom synthesized since they are project dependent, and thus are more expensive than commercial UCE probe sets.

Individual probes cost between $200 and $50 per sample depending on the quantity of probes purchased (see http://www.mycroarray.com/mybaits/mybaits-custom.html). Custom probes may also be prepared in the lab by attaching probes (commercially synthesized primers) to DynaBeads (ThermoFisher Scientific) via a biotion-streptavidin bond. The initial layout will be thousands of dollars with the actual cost largely determined by the number of synthesized probes, but thus prepared in-house probes will be sufficient for enriching thousands of individuals.

A summary is provided in Table 1.

**Table 1.** Comparison of hardware and reagent requirements to perform each of the four examined protocols.^1^ ^2^ ^3^

| Equipment | RAD | ddRAD | UCE | Exon-cap |
| --- | --- | --- | --- | --- |
| Sonicator/nebulizer/fragmentase | yes<sup>1</sup> | no | yes | yes |
| PippinPrep | optional | optional<sup>2</sup> | optional | optional |
| Basic PCR reagents | yes | yes | yes | yes |
| Restriction enzyme(s) | yes | yes | no | no |
| Hybridization probes | no | no | yes | yes |
| Barcoded adaptors | yes | yes | yes | yes |
| T4 DNA polymerase | yes | no | yes | yes |
| T4 DNA ligase | yes | yes | yes | yes |
| 3' → 5' proofreading DNA polymerase | yes | yes | yes | yes |
| Agencourt beads | yes | yes | yes | yes |
| Access to NGS machine | yes | yes<sup>3</sup> | yes | yes |

<sup>1</sup>in theory fragmentase could be used, but we do not know of its use in practice
<sup>2</sup>but recommended for multi-run experiments
<sup>3</sup>appropriate for an IONTORRENT PGM

### RADseq

The RADseq protocol requires an initial layout of several hundred dollars for the purchase of restriction and modifying enzymes, and several thousand dollars into the purchase of phosphorylated barcoded primers for linker construction. However, once purchased, enzymes can be used to prepare hundreds or thousands of samples, and primers can be used to prepare thousands of samples. Substantial savings can be made via the use of combinatorial barcoding (both ends of fragments are labelled with barcoded primers).

This protocol also requires access to a DNA sonicator or shearer, or fragmenting enzyme, and the sheared fragments will need to be polished before the second adapter can be added; the first adaptor is added at the restriction site. After sonication, all fragments are represented in any given size range and thus all RAD fragments are sequenced in a sequencing run. On an Illumina HiSeq 2500, at least 100 individuals can be sequenced at a minimum 10× read depth.

Floragenex (http://www.floragenex.com) provides a commercial service at approximately $100 per sample which includes library preparation and sequencing.

### ddRADseq

This protocol has the same requirements as RADseq with the difference that two restriction enzymes are used, and adaptors are added at the two restriction sites, and thus also no blunting and polishing step is required. The ddRADseq protocol is universally applicable for the same reason as the RADseq protocol. The use of the two restriction enzymes in the ddRADseq serves to generate in a reproducible number of fragments of different sizes. From this distribution of size fragments, a range is then chosen for sequencing, which in effect reduces the total number of unique fragments. In contrast to the RADseq protocol where all fragments are present in any give size range, in the ddRADseq protocol any given range of fragments has different combination and number of unique fragments. It is therefore critical that size selection for sequencing is replicable, and the same size range of fragments is selected between sequencing runs if sequencing runs are to be combined. Otherwise the total number of overlapping loci between sequencing runs will be low, hampering joint analyses. We therefore recommend the use of a PippinPrep or BluePippin for size selection.

The size selection normally results in the reduction of total number of unique fragments in a ddRADseq analysis to 1 /10th of the number of unique fragments expected in a RADseq analysis, which on Illumina HiSeq 2500 would allow the sequencing of 1,000 to 2,000 individuals at the same depth of coverage as in a RADseq experiment.

The relatively small number of fragments generated by this technique—generally between 5,000 and 15,000—makes the ddRADseq protocol one of the few protocols suitable for adaptation to the IonTorrent PGM, a machine that generates approximately 5 million sequencing reads, allowing for 20 to 60 individuals (taxon dependent), to be sequenced simultaneously at at least 10× read depth.

Again, Floragenex (http://www.floragenex.com) provides a commercial service at approximately $100 per sample which includes library preparation and sequencing.

### UCE

The UCE protocol requires investment into a sonicator/nebulizer, and the purchase of hybridization probes for each sample analyzed. Cost of a commercial probes sold by MyBaits ranges from $75 per sample (sufficient for 8 samples) to $30 per sample (sufficient for 96 samples) (see http://www.mycroarray.com/mybaits/mybaits-UCEs.html).

In theory the UCE protocol is universally or nearly universally applicable. While different probe sets have been developed for tetrapods and fishes, due to the ultraconserved nature of the probes, the same probes can be used in highly divergent taxa. The flanking regions contain the majority of the phylogenetic information, with flanking regions up to 200–400 bp from the UCE being alignable. Variation increases exponentially with distance from the core UCE region (Faircloth *et al*., 2012).

Enrichment of the total genomic DNA is on the order of 3.5% to 10% (Jake Enk, MycroArray Inc. pers. comm.; Smith *et al*., 2014) which means that 90–96% of the usable sequences from an NGS run are discarded. However, there are only 2,560 baits targeting 2,386 UCEs, and assuming that each UCE-containing contig is ≈ 600 bp long (160 bp UCE core plus flanking regions), approximately one hundred 100 bp reads per UCE will need to be assembled into a contig at an average 10× read depth. In the end, the number of individuals that can be analyzed per sequencing run is about half of that of a RADseq run. It should also be noted that the per-site phylogenetic information content of a UCE is about 1/3 that of a RADseq locus at the point of their maximal phylogenetic informativeness, and UCE loci are also contain much less phylogenetic information for very recent divergences.

RapidGenomics (http://www.rapid-genomics.com) provides a commercial service at approximately $160 per sample which includes library preparation and sequencing.

### Exon-cap

For exon-cap, whole genomic DNA has to be enriched for the targeted regions in the same manner as UCEs. Each project/taxonomic group will require the development of a specific probe set, although published probesets (e.g. Lemmon *et al*., 2012) are quite broad in their scope (e.g. vertebrates). There can be therefore a substantial bioinformatics investment in probe design for new groups (Bragg *et al*., 2015; Li *et al*., 2013), as well as the cost of the purchase of hybridization probes. However, a recent study by Li *et al*. (2013) demonstrated that it is possible to effectively capture orthologous regions in phylogenetically deeply divergent taxa using modified capture protocols. Custom probes for capture are sold commercially by MyBaits at a cost of $200 per sample (sufficient for exon capture in 12 samples) to $50 per sample (sufficient for 768 samples) (see http://www.mycroarray.com/mybaits/mybaits-custom.html). The 512-locus exon-cap probe-set represents one fifth of the number of loci of the UCE protocol. Assuming each exon locus is on average the same length as the UCE locus, five times as many individuals could be analyzed in a sequencing run when compared to the UCE or RADseq protocol.

No commercial service is currently available, but a collaborative project with the Lemmon laboratory at the University of Florida (http://anchoredphylogeny.com/) runs at around $12,000 per project analyzing approximate 100 samples.

### Sanger sequencing

By far the most expensive and time consuming technique is traditional Sanger sequencing. Obtaining data for the 512 loci targeted in the exon-cap protocol would require at least 1,000 sequencing reactions per individual (assuming each locus is approximately 700 bp in length). Collecting data for 100 individuals would thus require generating at least 100,000 sequences which is prohibitive both in cost and in time investment. For example Polina Perelman (pers. com.) estimated it took her over one year of continuous labwork to generate sequence data from 54 genes for 191 taxa (see Perelman *et al*., 2011). This represents one fifth the effort that would have been necessary to generate sequence data from the 512 exon-cap loci for 100 taxa if it were generated using traditional Sanger sequencing methods.

### Conclusions

The different genomic data types are effectively equivalent in their ability to robustly test phylogenetic hypotheses at the range of phylogenetic divergences analyzed in this study. Therefore what should guide a research labs choice of a technique should be other considerations that include access to equipment, laboratory expertise, data interchangeability, envisioned multiple use of data, etc.

A summary is provided in Table 2.

**Table 2.** Comparison the four examined protocols—attributes which may be considered when choosing a protocol.

| Attribute | Best | Worst |
| --- | --- | --- |
| Cost per sample | RAD/ddRAD | Exon-cap |
| Cost per bp | RAD/ddRAD | Exon-cap |
| Maximal per site phylogenetic informativeness | RAD | UCE |
| Total phylogenetic informativeness | RAD | Exon-cap |
| Phylogenetic informativeness at deep divergences | Exon-cap | ddRAD |
| Broadest phylogenetic/phylogeography suitability | RAD/ddRAD | Exon-cap/UCE |
| Resolving phylogenetically difficult problems | All other | ddRAD |
| Across laboratory dataset combinability | Exon-cap/UCE | ddRAD |
| Combinability with existing Sanger datasets | Exon-cap | All other |
| Suitability for low capacity NGS machine | ddRAD | All other |

## Footnotes

1 will be made available upon formal publication of the article

2 placed by some authors into the genus *Carlito*

1 in theory fragmentase could be used, but we do not know of its use in practice

2 but recommended for multi-run experiments

3 ppropriate for an IonTorrent PGM

